# Lack of p38 activation in T cells increases IL-35 production and protects against obesity by promoting thermogenesis

**DOI:** 10.1101/2023.08.04.551982

**Authors:** Ivana Nikolic, Irene Ruiz-Garrido, María Crespo, Rafael Romero-Becerra, Luis Leiva-Vega, Alfonso Mora, Marta León, Elena Rodríguez, Magdalena Leiva, Ana Belén Plata-Gómez, Maria Beatriz Alvarez, Jorge L. Torres, Lourdes Hernández-Cosido, Juan Antonio López, Jesús Vázquez, Alejo Efeyan, Pilar Martin, Miguel Marcos, Guadalupe Sabio

## Abstract

Obesity is characterized with low grade inflammation, energy imbalance and impaired thermogenesis. The role of regulatory T cells (Treg) in inflammation-mediated maladaptive thermogenesis has not been well established. We discovered that p38 pathway is a key regulator of T cell-mediated adipose tissue (AT) inflammation and browning. Mice with T cells specific deletion of the p38 activators, MKK3/6, were protected against diet-induced obesity and AT inflammation improving their metabolic profile, higher browning and thermogenesis. We identified IL-35 as a driver of adipocyte thermogenic program through ATF2/UCP1/FGF21 pathway. IL-35 limits CD8^+^ T cell infiltration and inflammation in AT. Interestingly, we found that IL35 was reduced in visceral fat from obese patients. Mechanistically we showed that p38 controls the expression of IL-35 in human and mouse Treg cells through mTOR pathway activation. Our findings highlight p38 signaling as a molecular orchestrator of AT T cell accumulation and function and identify p38 and IL-35 as promising targets for metabolic diseases.

## INTRODUCTION

Obesity is a serious worldwide health epidemic associated with an increased risk of life-threatening diseases (type 2 diabetes (T2D), cardiovascular diseases, and cancer) (1). The hallmark of obesity is a chronic low-grade inflammation, both systemically and in adipose tissue (AT), and this inflammation is one of the main promoters of insulin resistance and the development of T2D (2). Inflammatory myeloid cells, including macrophages and neutrophils, are key players in obesity comorbidities like diabetes and steatosis (3, 4). Chronic inflammation in obese AT is also mediated by adaptive immune cells, such as CD8^+^ and CD4^+^ T cells (5), and absence of these conventional T cells in mice reduces AT inflammation (6). CD8^+^ T cell infiltration and accumulation in AT plays a crucial role in the recruitment of macrophages and the maintenance of inflammation (7). Single cell RNA sequencing (scRNA-seq) of AT obtained from obese patients identified a potentially dysfunctional population of CD8^+^ T cells associated with metabolic disease (8). Moreover, whereas infiltration of AT by conventional T cells increases during obesity, numbers of protective Treg cells are sharply decreased, and expansion of this population ameliorates AT inflammation and insulin resistance (9, 10). Tregs are abundant in lean AT from mice and humans, and levels of the Treg hallmark transcription factor Foxp3 show an inverse correlation with body mass index (11), suggesting that preserving Treg accumulation in AT could be a novel anti-obesogenic strategy.

In addition to control the immune response within AT, immune cells also play a significant part in regulating AT thermogenic function. The recruitment of anti-inflammatory macrophages and type 2 innate lymphoid cells actively promotes adaptive thermogenesis (12, 13), while proinflammatory macrophages and CD8^+^ T cell are one of the main promoters of this compromised thermogenic capacity of AT (14, 15). However, the specific role of regulatory T (Treg) cells in thermogenesis remains unclear with data showing that this cell population facilitates obesity development and represses AT browning (16) as well as studies showing the protective role of Treg cells in promoting adaptive thermogenesis and browning (17, 18). Further research in this field is necessary to gain a comprehensive understanding of how Treg cells influence thermogenic mechanisms, which could provide valuable insights into the development of obesity and potential therapeutic strategies targeting AT metabolism.

Comprised of four members (p38α, p38β, p38γ, and p38δ), the p38 kinases belong to the stress protein kinase family and are activated by the upstream MAP kinase kinases MKK3 and MKK6 (19). This p38 pathway serves as a central regulator of the immune response and plays a critical role in the development of inflammatory diseases associated with obesity and becomes activated in various organs, including immune cells (1). For instance, the activation of p38γ/δ in neutrophils and p38α in macrophages promotes liver steatosis (3, 20). Interestingly, Treg cells present marked activation of the p38 pathway compared with conventional T cells (21), and lack of p38α/β in T cells enhances Treg cell induction in mice (22). Therefore, we wondered how p38 activation in T cells would affect AT inflammation and function during obesity. To address this question, we generated mice with specific deletion of the p38 family upstream activators MKK3 and MKK6 in conventional T cells (MKK3/6^CD4-KO^ mice). In our study we observed that lack of p38 activators results in increased population of Treg cells in circulation, lymph nodes and adipose tissue in obesity. Furthermore, we described a novel function of p38 kinases in Treg cells repressing IL-35 expression in human and mouse Treg cells through mTOR pathway inhibition. This Treg specific cytokine, which was also reduced in visceral fat from obese patients, restrains CD8^+^ T cell infiltration and inflammation in AT and induces adipocyte thermogenic program by activating ATF2 pathway leading to higher UCP1 and FGF21 levels. As consequence, mice lacking the upstream activators of p38s in T cells were protected against diet-induced obesity (DIO) and AT inflammation along with an improved metabolic profile characterized by enhanced browning and reduced liver steatosis. Our findings highlight the role of p38 signaling in reducing Treg accumulation in AT and identifies this kinase family as a key regulator in T cell function during obesity. In addition, we identify IL-35 as a potential therapeutic target for immunotherapy aimed at combating obesity.

## RESULTS

### 1.1. Lack of MKK3/6 in T cells protects against HFD-induced obesity, type II diabetes and liver steatosis

The significance of the p38 kinase family in innate cells, such as macrophages and neutrophils, in relation to obesity and metabolic diseases, has been extensively researched (1). However, the role of this stress kinase family in T cells during obesity is not fully understood. As an initial step, we took advantage of the data available from single cell RNA sequencing of human AT from White Adipose Atlas (23) and checked the expression pattern of the p38 signaling pathway in the cluster of T cells infiltrating AT from lean individuals (BMI <30) and patients with obesity (BMI 30-40) or severe obesity (BMI 40-50) (Fig. 1A and table S1). Patients with severe obesity (BMI 40-50) presented a signature of activation of p38 pathway in T cells from AT, compared with lean individuals (BMI 20-30). Precisely, we found higher expression of p38α (*MAPK14*), its up-stream activator, MKK6 (*MAP2K6)* and its substrate *ATF2* and down-expression of the negative regulator the phosphatase *DUSP1* (Fig. 1B and 1C). All together, these data suggest that p38 signaling pathway have a role in T cells during obesity.

**Fig. 1.**
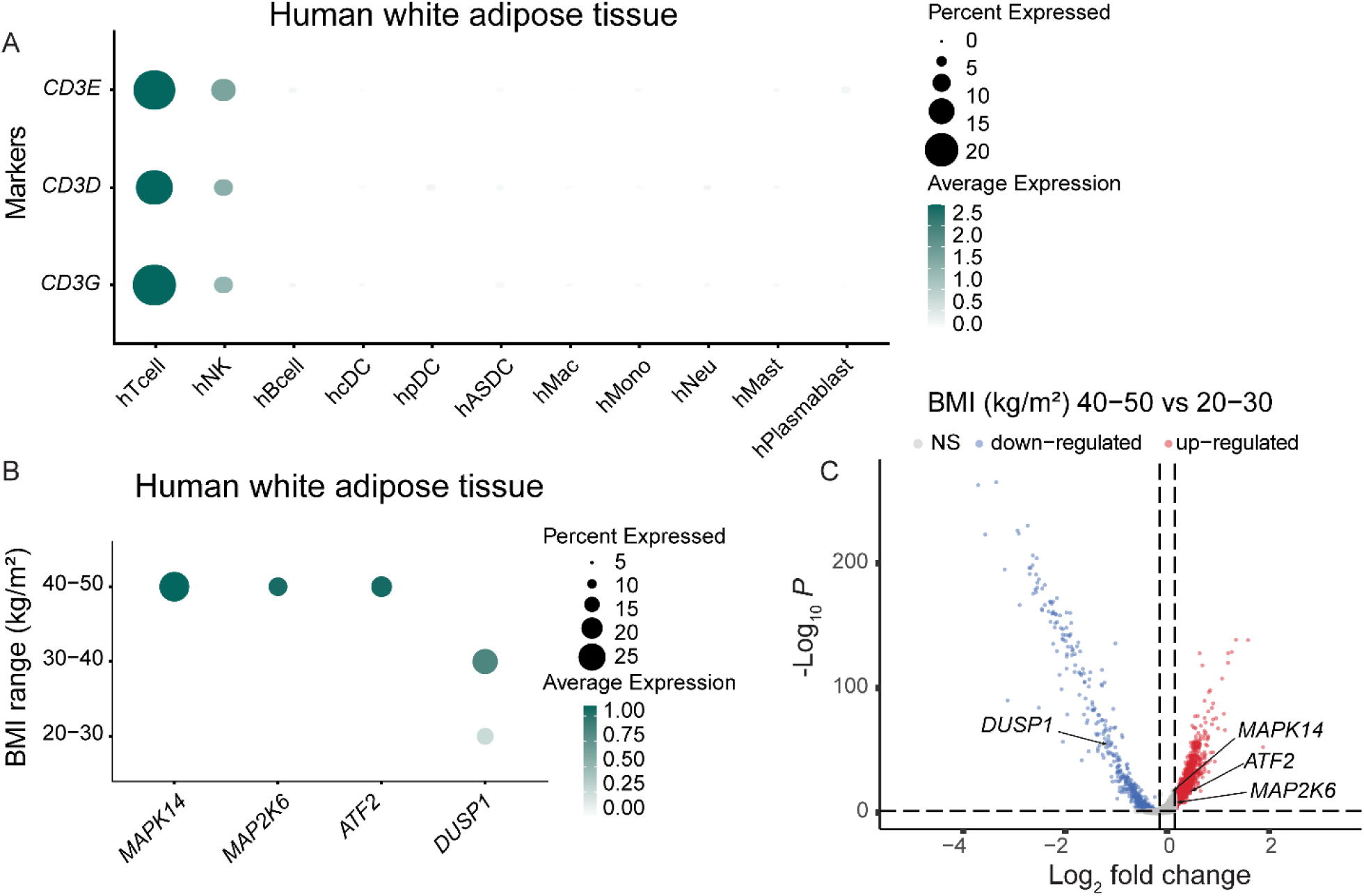
p38 MAPK pathway is upregulated in T cells in obese human white adipose tissue. (**A**) Dot plot of the expression of the indicated T cell marker genes Hin the different cell type clusters in human white adipose tissue single-cell RNA-seq data from Emont et al. (18). (**B**) Dot plot of the expression of the indicated genes by BMI range in human white adipose tissue T cell cluster shown in (A). (**C**) Volcano plot of differentially expressed genes in human white adipose tissue T cell cluster in severe obese (BMI 40-50 kg/m2) versus non-obese (BMI 20-30 kg/m2) subjects. The vertical dashed line indicates a log2 fold change cut-off of 0.15. The horizontal dashed line indicates a -log10 adjusted p-value (using Bonferroni correction) cut-off of 1.3 (adj p-value < 0.05). DC: dendritic cells; Mac: macrophages; Mono: monocytes; Neu: neutrophils; Mast: mastocytes.

To evaluate this hypothesis, we generated mice lacking the p38 upstream activators, MKK3 and MKK6, specifically in conventional T cells (MKK3/6^CD4-KO^). Specific deletion of both kinases was observed in CD4^+^ and CD8^+^ cells, whereas expression was unaffected in other cell types (NK, liver and AT) (Fig. S1A). Since previous studies using mice lacking p38 family members (22, 24) showed lymphoid dystrophy we evaluated whether lack of p38 activation leads to lymphocyte abnormalities. Analysis of thymus, spleen and periphery lymphoid organs weight and cell number showed no differences in MKK3/6^CD4-KO^ mice compared to control mice CD4-Cre (Fig. S1B and S1C). In addition, we didn’t observe differences in T cell development in thymus (Fig. S1D). Furthermore, we measured real-time bioenergetic changes naïve CD4^+^ T cells activated with PMA/Ionomycin and found no differences between genotypes (Fig. S1E). These data suggest that in contrast with deficiency in p38s, lack of p38 activation in T cells does not induces lymphoid dystrophy nor problems in T cell activation.

Interestingly, MKK3/6^CD4-KO^ mice were leaner than control CD4-Cre mice and had lower epididymal WAT (eWAT) and subcutaneous WAT (sWAT) weight (Fig. S1F and S1G). Metabolic cages analysis revealed increased energy expenditure (EE) in chow-diet fed MKK3/6^CD4-KO^ mice (Fig. S2A), without differences in food intake or locomotor activity (Fig. S2B). In addition, chow-diet fed MKK3/6^CD4-KO^ mice presented elevated brown adipose tissue (BAT) thermogenesis to judge by higher interscapular temperature (Fig. S2C-D) in MKK3/6^CD4-KO^ mice compared either to CD4-Cre or littermates (MKK3/6^f/f^). These data suggest that p38 activation in T cells regulates BAT thermogenesis and energy homeostasis and might have an important role in the development of obesity-associated metabolic diseases.

To further elucidate the contribution of p38 activation in T cells during obesity, we fed animals a high-fat diet (HFD) for 8 weeks. MKK3/6^CD4-KO^ mice were protected against HFD-induced obesity (Fig. 2A). The lower body-weight correlated with lower body and fat mass measured by MRI (Fig. 2B), reduced fat-depot and liver mass (Fig. 2C). Similar phenotype was observed in MKK3/6^CD4-KO^ female mice (Fig. S3A-S3C). Since obesity is usually followed by hyperglycemia and ectopic fat accumulation in liver (also known as liver steatosis) (1), analysis of blood glucose and liver histology showed protection against hyperglycemia and steatosis in HFD-fed MKK3/6^CD4-KO^ mice (Fig. 2D and 2E). We also performed glucose and insulin tolerance test (GTT and ITT), and found no differences in glucose tolerance and significantly improved insulin sensitivity in HFD-fed MKK3/6^CD4-KO^ mice (Fig. S4).

**Fig. 2.**
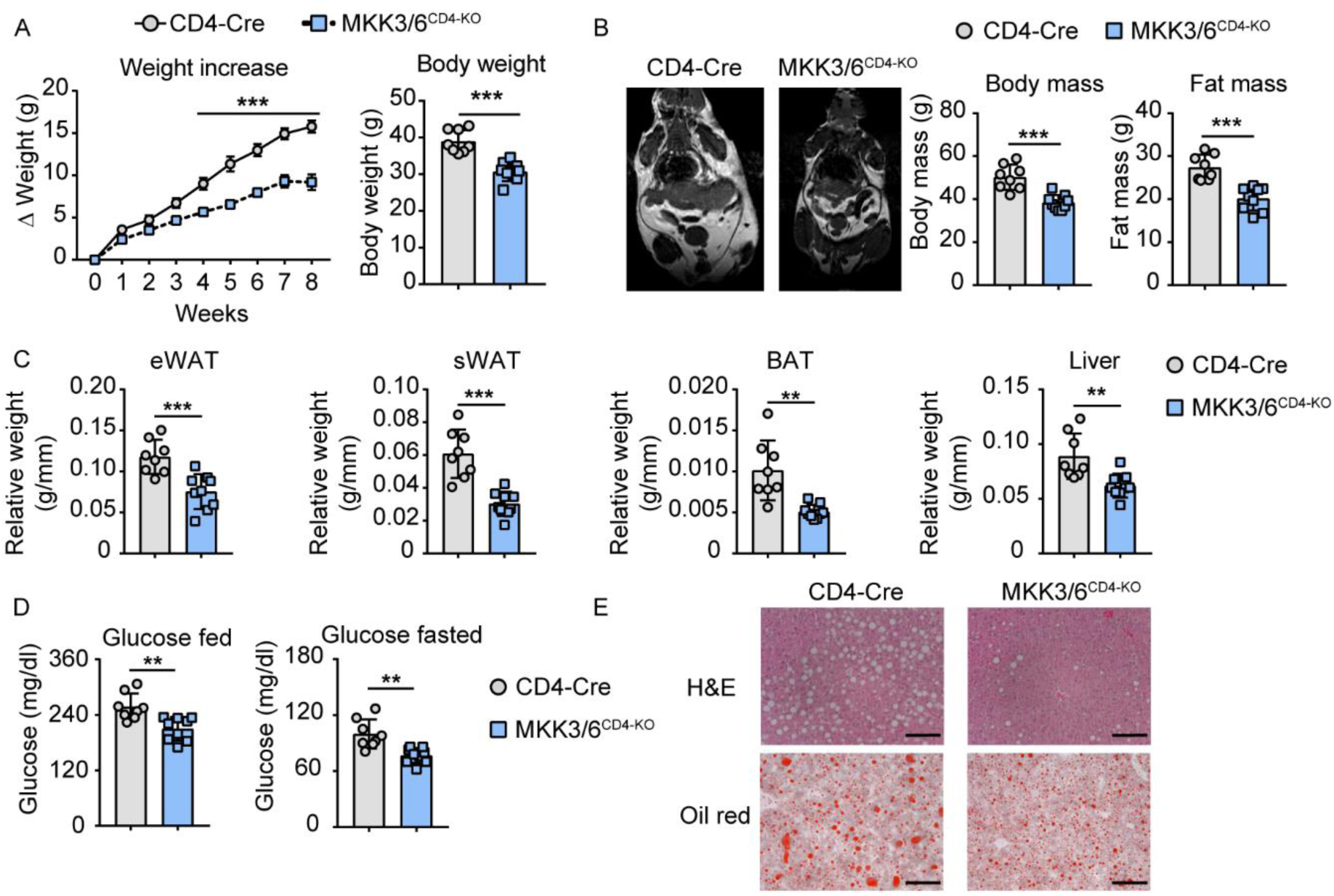
MKK3/6 deficiency in T cells protects against HFD-induced obesity. MKK3/6^CD4-KO^ and CD4-Cre mice were fed a high-fat diet (HFD) for 8 weeks. (**A**) Body weight evolution in CD4-Cre and MKK3/6^CD4-KO^ male mice fed the HFD for 8 weeks (starting at 8-10 weeks old). Data are presented as the increase above initial weight (left) and absolute weight at the end of the experiment (right) (mean ± SEM; CD4-Cre n = 8 mice; MKK3/6^CD4-KO^ n = 9 mice). (**B**) NMR analysis of body and fat mass in MKK3/6^CD4-KO^ and CD4-Cre mice after 8 weeks of HFD (mean ± SEM; CD4-Cre n = 8 mice; MKK3/6^CD4-KO^ n = 9 mice). Representative images are shown on the left. (**C**) eWAT, sWAT, BAT, and liver mass relative to tibia length (mean ± SEM; CD4-Cre n = 8 mice; MKK3/6^CD4-KO^ n = 9 mice). (**D**) Blood glucose concentration after 8 weeks of HFD in mice fed (left) or fasted overnight (right) (mean ± SEM; CD4-Cre n = 8 mice; MKK3/6^CD4-KO^ n = 9 mice). (**E**) Representative haematoxylin–eosin and oil-red O staining of liver sections. Scale bar, 100 µm. Data are mean ± SEM, *p < 0.05, ** p < 0.01, ***p < 0.001 CD4-Cre versus MKK3/6^CD4-KO^. Analysis by 2-way ANOVA coupled to the Bonferroni post-test (A) or *t* test or by the Welch test when variances were different (A-D).

### 1.2. Deficiency of MKK3/6 in T cells increases AT thermogenesis and browning

Immune system has been shown to regulate body weight through the regulation of AT thermogenesis (25). Therefore, we measured BAT temperature, and observed HFD-fed MKK3/6^CD4-KO^ mice (both male and female) had higher skin temperature surrounding interscapular BAT (Fig. 3A and S3D), suggesting that lack of p38 activation in T cells leads to higher BAT thermogenesis. To confirm that these differences are also present in thermoneutrality conditions, we placed CD4-Cre and MKK3/6^CD4-KO^ mice at 30 °C during the whole course of HFD. We observed that at isothermal housing MKK3/6^CD4-KO^ mice maintained their protection against HFD-induced obesity (Fig. 4A), reduced body and fat mass evaluated by MRI (Fig. 4B), reduced fat depots (Fig. 4C), adiposity (Fig. 4D) and liver steatosis (Fig. 4D). In addition, MKK3/6^CD4-KO^ mice preserved higher BAT temperature at thermoneutrality (Fig. 4E), confirming that protection against obesity is autonomous and independent of animal housing temperature.

**Fig. 3.**
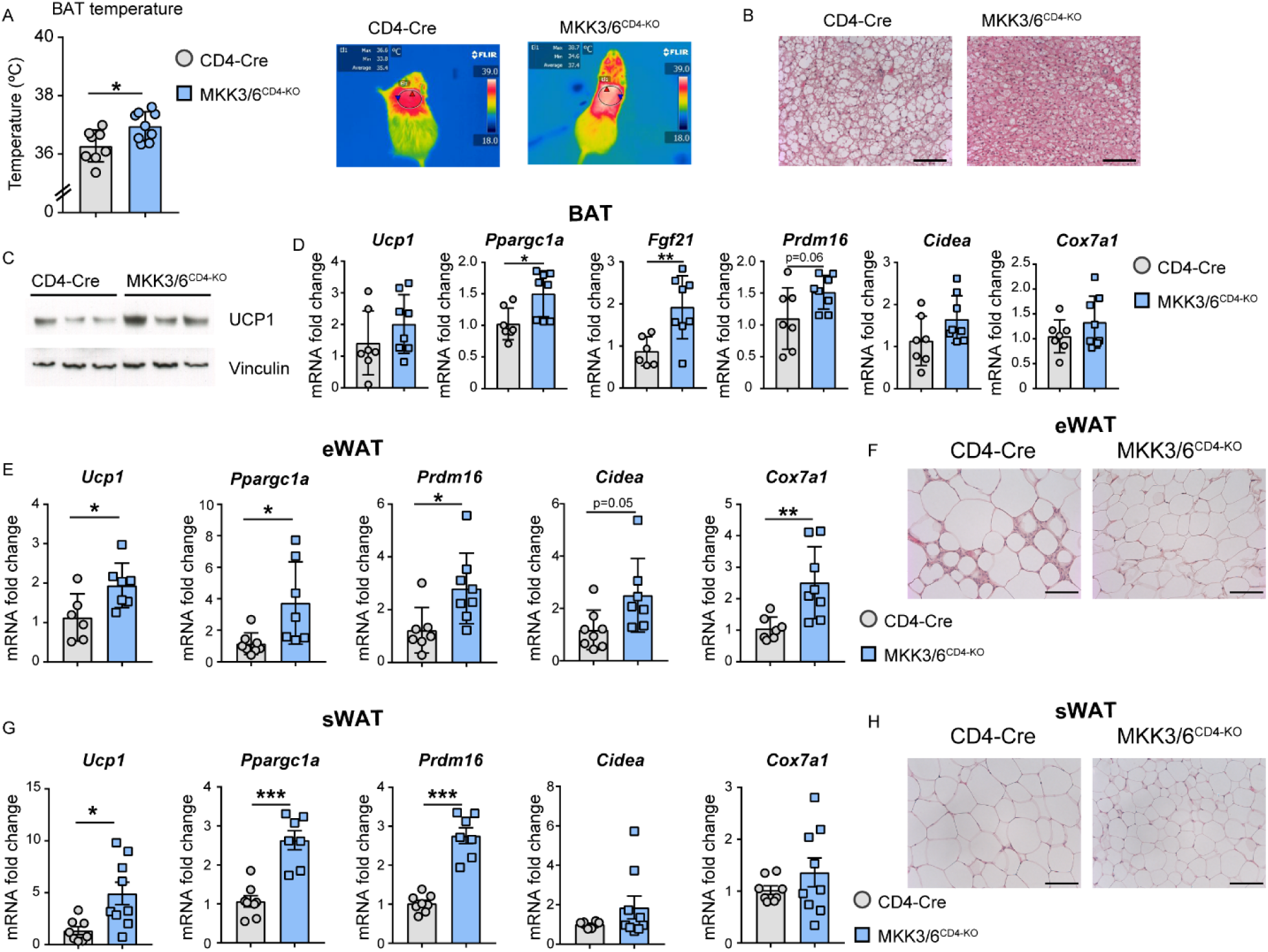
Lack of MKK3/6 in T cells increases BAT thermogenesis and AT browning. MKK3/6^CD4-KO^ and control (CD4-Cre) mice were HFD-fed for 8 weeks. (**A**) Skin temperature surrounding interscapular BAT. Right panels show representative infrared thermal images (mean ± SEM; CD4-Cre n = 8 mice; MKK3/6^CD4-KO^ n = 9 mice). (**B**) Representative H&E staining of BAT sections. Scale bar, 100 µm. (**C**) Western blot analysis of UCP1 in BAT. (**D**) qRT-PCR analysis of thermogenic gene mRNA expression in BAT isolated from control or MKK3/6^CD4-KO^ mice. mRNA expression was normalized to the expression of β-actin mRNA and presented as fold increase compared to CD4-Cre (mean ± SEM; CD4-Cre n = 7 mice; MKK3/6^CD4-KO^ n = 8 mice). (**E**) qRT-PCR analysis of browning genes mRNA expression from eWAT isolated from control or MKK3/6^CD4-KO^ mice. mRNA expression was normalized to β-actin mRNA and presented as fold increase compared to CD4-Cre (mean ± SEM; CD4-Cre n = 7 mice; MKK3/6^CD4-KO^ n = 8 mice). (**F**) Representative H&E staining of eWAT sections. Scale bar, 100 µm. (**G**) qRT-PCR analysis of browning genes mRNA expression from sWAT isolated from control or MKK3/6^CD4-KO^ mice. mRNA expression was normalized to β-actin mRNA and presented as fold increase compared to CD4-Cre (mean ± SEM; CD4-Cre n = 7 mice; MKK3/6^CD4-KO^ n = 8 mice). (**H**) Representative H&E staining of sWAT sections. Scale bar, 100 µm. Data are mean ± SEM, *p < 0.05, ** p < 0.01, ***p < 0.001 CD4-Cre versus MKK3/6^CD4-KO^. Analysis by t test or by the Welch test when variances were different (A, B, D, F, H).

**Fig. 4.**
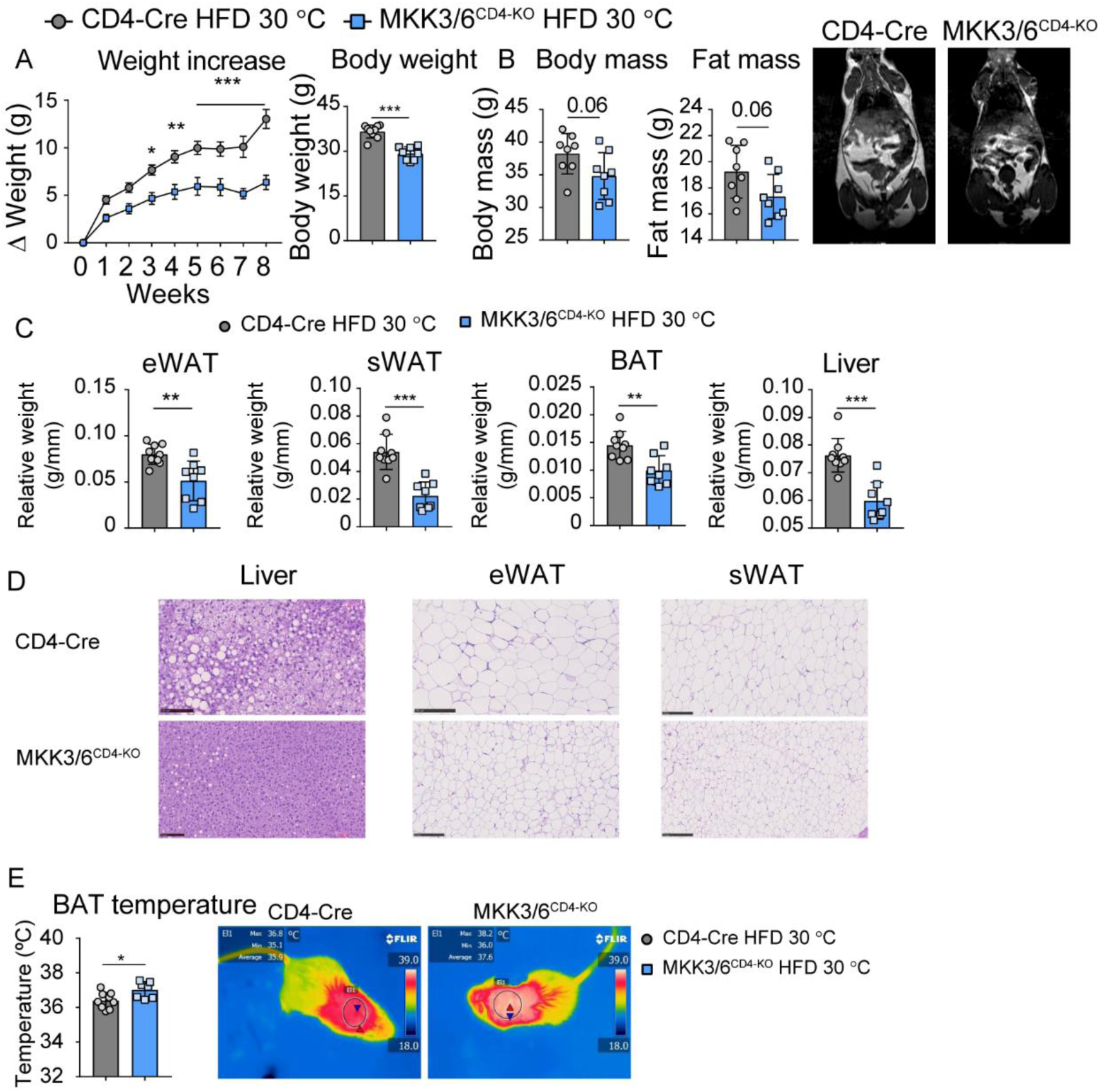
MKK3/6 deficiency in T cells protects against HFD-induced obesity in isothermal housing. MKK3/6^CD4-KO^ and CD4-Cre mice were fed a high-fat diet (HFD) for 8 weeks and housed at 30 °C during whole course of HFD. (**A**) Body weight evolution in CD4-Cre and MKK3/6^CD4-KO^ male mice fed the HFD for 8 weeks (starting at 8-10 weeks old). Data are presented as the increase above initial weight (left) and absolute weight at the end of the experiment (right) (mean ± SEM; CD4-Cre n = 9 mice; MKK3/6^CD4-KO^ n = 8 mice). (**B**) NMR analysis of body and fat mass in MKK3/6^CD4-KO^ and CD4-Cre mice after 8 weeks of HFD (mean ± SEM; CD4-Cre n = 9 mice; MKK3/6^CD4-KO^ n = 8 mice). Representative images are shown on the right. (**C**) eWAT, sWAT, BAT, and liver mass relative to tibia length (mean ± SEM; CD4-Cre n = 9 mice; MKK3/6^CD4-KO^ n = 8 mice). (**D**) Representative haematoxylin– eosin and oil-red O staining of liver, eWAT and sWAT sections. Scale bar, 100 µm for liver and 250 µm for eWAT and sWAT. (**E**) Skin temperature surrounding interscapular BAT. Right panels show representative infrared thermal images (mean ± SEM; CD4-Cre n = 8 mice; MKK3/6^CD4-KO^ n = 7 mice). Data are mean ± SEM, *p < 0.05, ** p < 0.01, ***p < 0.001 CD4-Cre versus MKK3/6^CD4-KO^. Analysis by 2-way ANOVA coupled to the Bonferroni post-test (B) or *t* test or by the Welch test when variances were different (B, C, D).

Next, we analyzed BAT morphology and histological analysis revealed increased cellularity and preserved BAT morphology with typical multilocular lipid droplets (Fig. 3B). Multilocular cells with high energy-dispensing activity have been linked to elevated expression of the BAT marker mitochondrial UCP1 (26). UCP1 is a crucial mitochondrial protein involved in BAT thermogenesis which uncouples oxidative phosphorylation from ATP production, therefore dissipating chemical energy as heat (27). Quantitative RT-PCR and western blot analysis confirmed high UCP1 expression in BAT from HFD-fed MKK3/6^CD4-KO^ mice (Fig. 3C and 3D), correlating with the BAT activation and elevated thermogenesis-related genes *Fgf21* and its target *Ppargc1a* in HFD-fed MKK3/6^CD4-KO^ mice (Fig. 3C). It has been shown that FGF21 can act as an autocrine factor in adipocytes and induce UCP1 expression and thermogenic program (28). These results indicate that deficiency of p38 activation in T cells promotes BAT thermogenesis thus restraining obesity development.

T cells have been shown to regulate the browning of WAT (15, 29), as well as adipose derived FGF21 in autocrine/paracrine manner (30). Then we evaluated whether lack of p38 activation in T cells induced browning. We found significant upregulation of the browning-signature markers *Ucp1*, PR domain-containing16 (*Prdm16*), and peroxisome proliferator-activated receptor gamma (*Ppargc1*) in eWAT and subcutaneous WAT (sWAT) from HFD-fed MKK3/6^CD4-KO^ mice (Fig. 3E and 3G). We also observed smaller adipocyte size in both eWAT and sWAT from HFD-fed MKK3/6^CD4-KO^ mice (Fig. 3F and 3H). Given the association of obesity with impaired AT metabolism (26) and the emerging evidence for the role of T lymphocytes in the regulation of AT (31), we profiled metabolism-related gene expression in eWAT and sWAT from HFD-fed animals. RT-PCR revealed increased expression of genes involved in adipogenesis (*AdipoQ* and *Plin1*), lipogenesis (*Ppard, Acaca*, and *Scd1*), β-oxidation (*Acox1* and *Cpt1*), and glycolysis (*Pepck*) in HFD-fed MKK3/6^CD4-KO^ eWAT and sWAT (Fig. S5A and S5B). These results provide evidence for a T cell–AT crosstalk, regulated by p38 activation, that controls AT metabolism and browning, especially in obesity.

### 1.3. T cell MKK3/6-deficiency increases Treg accumulation and prevents CD8^+^ infiltration in AT

Obesity is considered as low-grade inflammation condition, so we next examined the cellular mechanism underlying the observed DIO protection by flow cytometry (gating strategy presented in Fig. S6). First, we analyzed CD4^+^, CD8^+^ and Treg population in spleen, blood and AT draining lymph node after HFD. While there were no differences in CD4^+^ and CD8^+^ cells in observed lymphoid compartments and circulation (Fig. S7A-S7C), we found more Treg cells in spleen, blood and lymph nodes in mice deficient for p38 activation in T cells after HFD (Fig S7A -S7C), consistent with previously published data showing that p38 deficiency enhances Treg induction (22). Obesity leads to decrease of protective Treg cells in AT (11), hence we wondered whether observed higher frequency of Treg cells in MKK3/6^CD4-KO^ mice also occurs in AT. Our results demonstrate that p38 activation in T cells inhibits Treg cell accumulation in AT and, in consequence HFD-fed MKK3/6^CD4-KO^ mice show AT Treg cell expansion (Fig. 5A). To corroborate this, we again checked whether p38 signalling pathway was differentially expressed in the cluster of Treg cells (Fig. S8A) in the White Adipose Atlas (23). We observed higher expression of p38α (*MAPK14*) in adipose tissue Treg cells from human with obesity (BMI 30-50) or with severe obesity (BMI 40-50) compared with lean individuals (BMI 20-30) (Fig. S8B, S8C, S8D and S8E and tables S2 and S3). We also found up-regulated its upstream activators MKK6 (*MAP2K6*, Fig. S6B) and MKK3 (*MAP2K3*) in Treg cells from human with obesity (Fig. S8B, S8F, S8G and table S4), and down-regulated expression of phosphatase *DUSP1* in Treg cells from human with severe obesity (Fig. S8B). In addition, *ATF-2* was also up-regulated in adipose tissue Treg cells from human with BMI 40-50 (Fig. S8B). All together, these data indicate that p38 signalling pathway is up-regulated in Treg cells from adipose tissue of human with obesity.

**Fig. 5.**
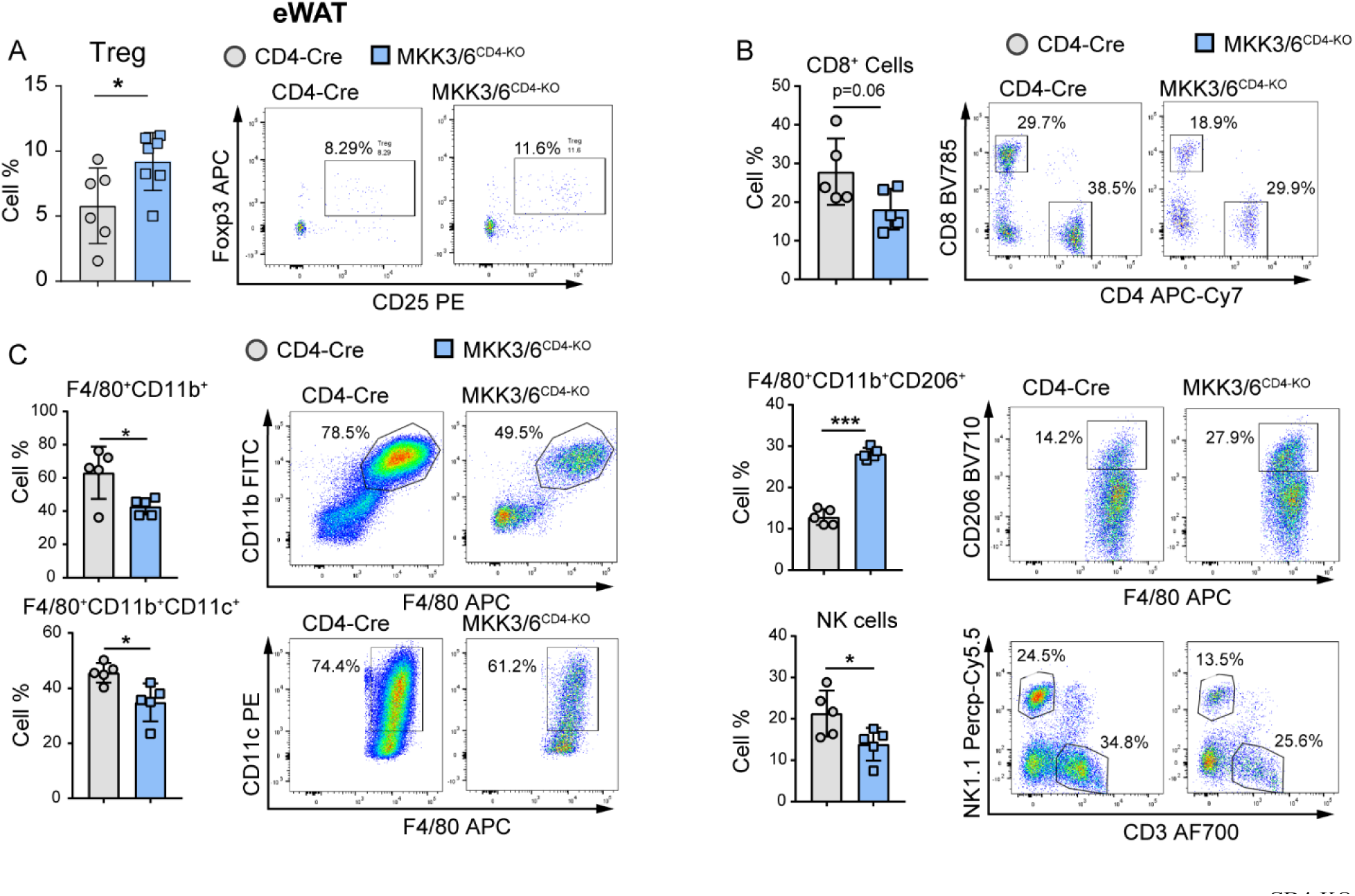
MKK3/6 deficiency in T cells promotes Treg cell accumulation in AT. MKK3/6^CD4-KO^ and CD4-Cre mice were fed a high-fat diet (HFD) for 8 weeks. (**A**) FACS quantification and representative dot plots of Treg cells (CD4^+^CD25^+^Foxp3^+^) in SVF (mean ± SEM; CD4-Cre n = 7 mice; MKK3/6^CD4-KO^ n = 7 mice) and (**B**) CD8^+^ T cells in SVF (mean ± SEM; CD4-Cre n = 5 mice; MKK3/6^CD4-KO^ n = 5 mice). (**C**) FACS quantification and representative dot plots of myeloid (F4/80^+^CD11b^+^), M1 macrophages (Mϕ) (F4/80^+^CD11b^+^CD11c^+^), M2 Mϕ (F4/80^+^CD11b^+^CD206^+^), and NK (NK1.1^+^CD3^-^) cells in SVF (mean ± SEM; CD4-Cre n = 5 mice; MKK3/6^CD4-KO^ n = 5 mice). Data are mean ± SEM, *p < 0.05, ** p < 0.01, ***p < 0.001 CD4-Cre versus MKK3/6^CD4-KO^. Analysis by *t* test or by the Welch test when variances were different (all graphs).

Since Treg cells possess immunosuppressive properties and may play a role in suppressing infiltration of CD8^+^ obesogenic T cell in AT (11, 32), we measured CD8^+^ T cells in eWAT by FACS. HFD-fed MKK3/6^CD4-KO^ mice have lower CD8^+^ cell infiltration in eWAT than HFD-fed CD4-Cre mice (Fig. 5B). CD8^+^ cell infiltration precedes macrophage accumulation and promotes proinflammatory macrophage recruitment (7), in concordance, eWAT from HFD-fed MKK3/6^CD4-KO^ mice presented less abundance of obesogenic, proinflammatory macrophages and NK cells while the frequency of anti-inflammatory macrophages (F4/80^+^CD11b^+^CD206^+^) was higher (Fig. 5C). These data suggest that p38 activation deficiency in T cells prevents AT inflammation.

### 1.4. p38 activation in Treg cells represses the expression of *Il-35* via mTOR pathway

In order to investigate the molecular mechanisms through which Treg cells control AT inflammation, we assessed the expression of IL-35, an immunosuppressive cytokine produced by Treg cells, in both the AT and stromal vascular fraction (SVF). Expression of Ebi3 and p35 the two subunits of the IL-35 were upregulated in HFD-fed MKK3/6^CD4-KO^ SVF (Fig. 6A and 6B), highlighting p38s as regulators of IL-35 production in Treg cells. To confirm this, we analyzed IL-35^+^ Treg cells in AT draining lymph nodes after HFD, and we observed higher frequency of IL35^+^ Treg cells in HFD-fed MKK3/6^CD4-KO^ mice (Fig. 6C). In order to gain deeper insights into the molecular mechanism underlying the control of IL35 production by p38 activation, we next measured *Il-35* expression in *in vitro* induced Treg cells (Fig. 6D). In line with the *in vivo* data from SVF and lymph nodes, expression of *Il-35* (p35) was higher in MKK3/6^CD4-KO^ Treg cells than in their CD4-Cre counterparts (Fig. 6D). To confirm the action of this pathway in human Treg cells, we inhibited p38 with the p38 pan inhibitor BIRB796 (33). As expected, p38 inhibition in human Treg cells induced the expression of *IL-35* (*P35*) (Fig. 6E).

**Fig. 6.**
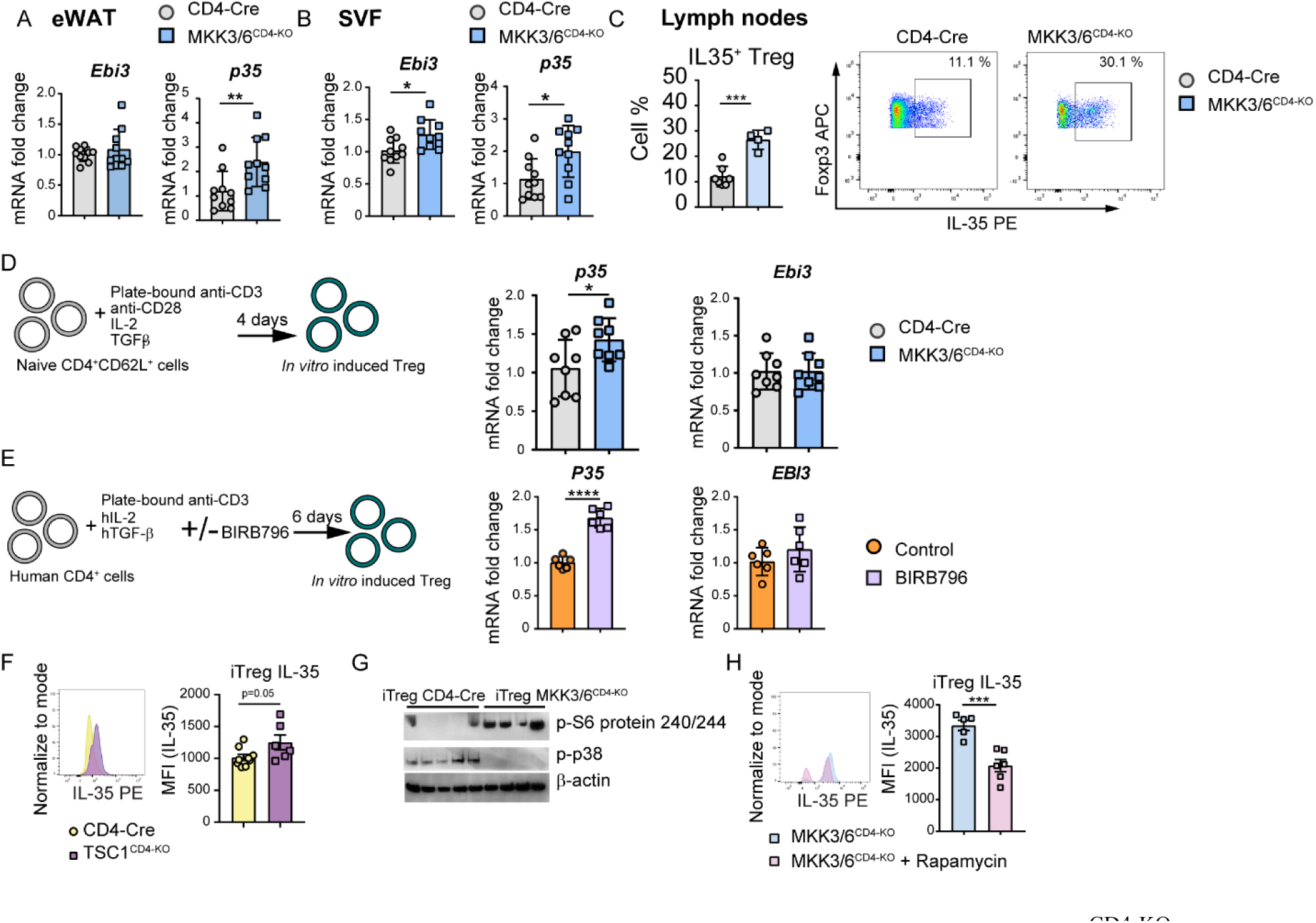
p38 activation in Treg cell inhibits IL-35 production. (**A-C**) MKK3/6^CD4-KO^ and CD4-Cre mice were fed a high-fat diet (HFD) for 8 weeks. (**A-B**) qRT-PCR analysis of mRNA expression in (**A**) eWAT and (**B**) SVF isolated from control or MKK3/6^CD4-KO^ mice. mRNA expression was normalized to *b-actin* mRNA (mean ± SEM; CD4-Cre n = 10 mice; MKK3/6^CD4-KO^ n = 10 mice). (**C**) FACS quantification and representative dot plots of IL-35^+^Treg cells in lymph nodes (mean ± SEM; CD4-Cre n = 7 mice; MKK3/6^CD4-KO^ n = 4 mice). (**D**) *In vitro* Treg cell induction (iTreg). Naïve CD4^+^ T cells were isolated from the spleens of CD4-Cre and MKK3/6^CD4-KO^ mice stimulated for 96h with plate-bound anti-CD3, soluble anti-CD28 + IL-2 +TGFβ. qRT-PCR analysis of *p35* and *Ebi3* mRNA in iTregs derived from control or MKK3/6^CD4-KO^ mice. mRNA expression was normalized to *b-actin* mRNA (mean ± SEM; CD4-Cre n = 8 wells; MKK3/6^CD4-KO^ n = 8 wells, technical replicates). (**E**) Induction of iTregs from CD4^+^ T cells isolated from healthy donor buffy coats and stimulated with plate-bound anti-CD3, soluble hIL-2 + hTGFβ for 6 days in the presence or absence of the p38 pan inhibitor BIRB796. qRT-PCR analysis of *P35* and *EBI3* mRNA in iTregs. mRNA expression was normalized to *GAPDH* mRNA (mean ± SEM; Control n = 6 wells; BIRB796 treated n = 6 wells, technical replicates). (**F**) FACS analysis of IL-35 MFI in *in vitro* induced iTreg cells from CD4-Cre and TSC1^CD4-KO^ mice (mean ± SEM; CD4-KO n = 9 wells; TSC1^CD4-KO^ n = 6 wells, biological replicates). (**G**) Western blot analysis of p-p70 T389, p-s6 protein S240/244 and p-p38 Thr180/Tyr182 in iTreg cells from CD4-Cre (n=5) and MKK3/6^CD4-KO^ mice (n=4). Loading control for p-p38 was run on different gel and not presented. (**H**) FACS analysis of IL-35 MFI in *in vitro* induced iTreg cells from MKK3/6^CD4-KO^ mice in the presence or absence of rapamycin for 4h (mean ± SEM; MKK3/6^CD4-KO^ n = 5 wells; MKK3/6^CD4-KO^ + rapamycin n = 6 wells, technical replicates). Data are mean ± SEM, *p < 0.05, ** p < 0.01, ***p < 0.001. Analysis by *t* test.

Previous studies have proposed a potential role of mTOR in regulating IL-35 levels (34). In concordance we found that Treg cells from mice with increased mTOR activation (TSC1^CD4-KO^ mice) presented higher levels of IL-35 (Fig. 6F). mTOR has been shown to be modulated by p38 (35). In agreement, western blot analysis demonstrated that lack of p38 activation in Treg cells leads to increase mTOR activation and higher S6 protein phosphorylation (Fig. 6G). To evaluate whether mTOR activation in MKK3/6^CD4-KO^ Treg cells was responsible for the increase in IL35 expression we inhibited mTOR pathway with rapamycin. We found that IL35 expression is dependent of mTOR activation since treating Treg cells lacking p38 activation (MKK3/6^CD4-KO^) with rapamycin diminished the IL35 expression (Fig. 6H). In summary, we described novel mechanism by which p38 kinases inhibits mTOR signaling pathway that in turn reduces IL-35 production in mouse and human Treg cells. Inhibition of p38s (genetically or chemically) resulted in increased IL-35 mRNA and protein levels.

### 1.5. Treg derived IL-35 triggers AT browning and is reduced in obese patients

As MKK3/6^CD4-KO^ mice expressed higher levels of IL-35 in AT, we postulated that IL-35 might have a role in AT thermogenesis and consequently in protection against obesity. Interestingly, we found a tendency to lower p35 expression in visceral adipose tissue obtained from obese patients compared to lean controls (Fig. 7A). Furthermore, MKK3/6^CD4-KO^ mice exposed to cold preserved their body and BAT temperature, while the temperature of control CD4-Cre mice gradually dropped during cold challenge (Fig. 7B). We next evaluated the functional role of IL-35 in thermogenesis *in vivo*. IL-35 injection triggers thermogenesis in mice compared with PBS treated group (Fig. 7C). To understand mechanistically how IL-35 cytokine controls thermogenesis in adipocytes, we treated brown adipocytes with IL-35 and found increased levels of UCP1 and FGF21 suggesting an activation of brown thermogenic program (Fig. 7D). IL-35 treatment leads to ATF-2 phosphorylation (Fig. 7E), the transcription factor that controls *Ucp1* and *Pgc1a* expression in BAT (36). This increased in *Ucp1* expression induced by IL-35 was down-regulated when ATF2 activation in adipocytes was blunted by using inhibitor of upstream activator of ATF2 (SB203580, p38α/β inhibitor) (Fig. 7F). Furthermore, IL35 also promoted adipogenesis in differentiated primary adipocytes which was reduced when ATF2 activation was suppressed (Fig. 7G). In brief, we described novel role of Treg derived cytokine IL-35 in controlling adipose tissue thermogenesis *via* pATF-2/UCP1/FGF21 axis.

**Fig. 7.**
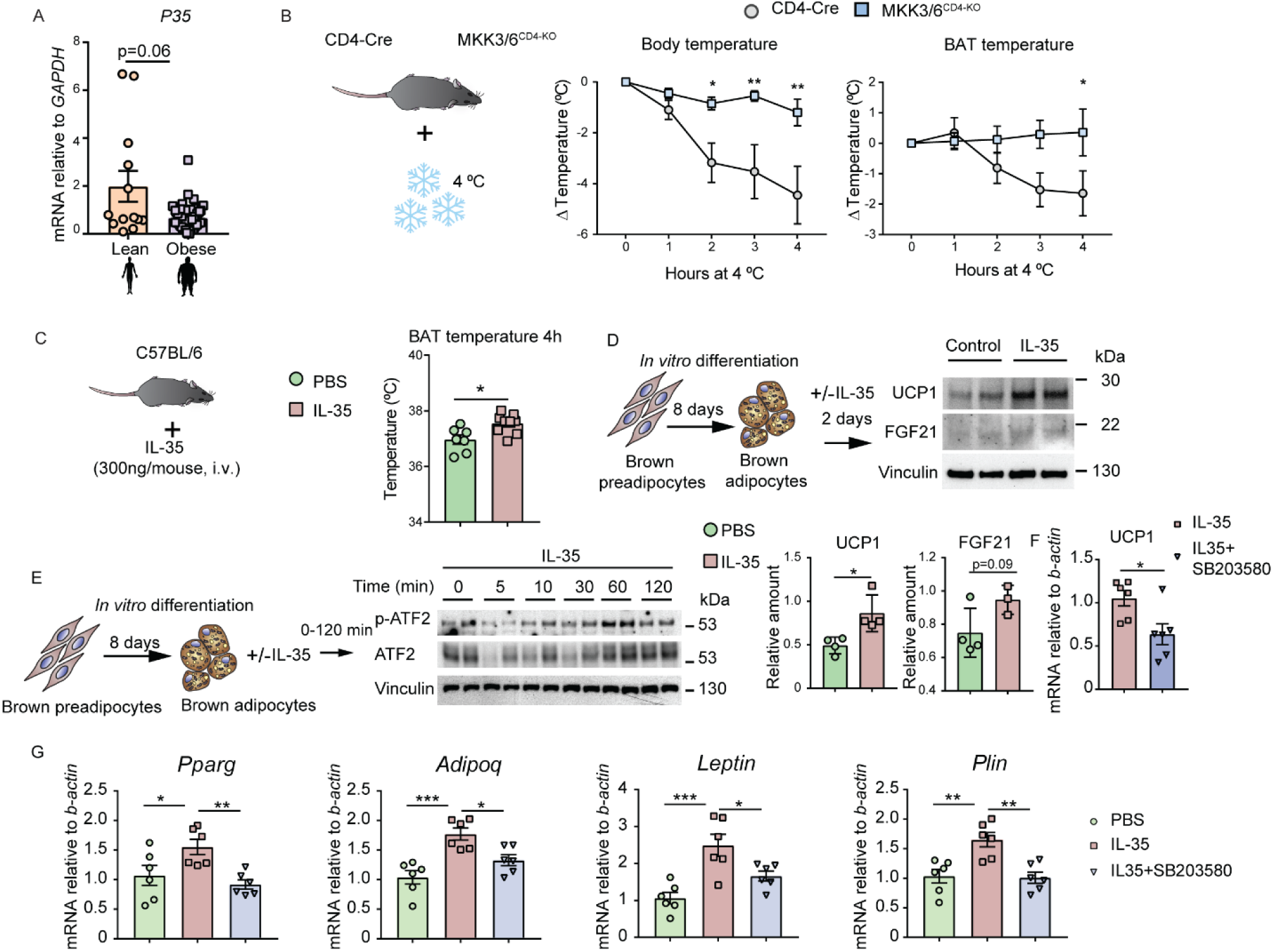
Treg-derived IL-35 promotes thermogenesis by increasing ATF-2 phosphorylation and UCP1 and FGF21 levels. (**A**) mRNA expression of p35 subunit of IL-35 in human visceral fat isolated from lean (N=12) and obese patients (N=52). (**C**) C57BL6 mice were treated with recombinant IL-35 i.v. (300ng per mouse) and BAT temperature was measured 4h later (mean ± SEM; PBS n = 7; IL-35 treated mice n = 9). (**D**) Immortalized brown preadipocytes were differentiated *in vitro*. Once differentiated, cells were stimulated in the presence or absence of IL-35 (100 ng/ml) for 48h and UCP1 and FGF21 levels were analysed by immunoblot. Loading control for UCP1 was run on different gel and not presented. (n=4 for each condition, technical replicates). (**E**) Immortalized brown preadipocytes were differentiated *in vitro*. Once differentiated, cells were stimulated in the presence or absence of IL-35 (100 ng/ml) for 0-120 minutes and ATF2 phosphorylation was analysed by immunoblot. (**F**) Differentiated adipocytes were stimulated with IL-35 (100 ng/ml) for 48h in the presence or absence of SB203580 inhibitor (10 uM). The expression of *Ucp1* level was measured by qRT-PCR and relativized to *b-actin*. (**G**) Primary white preadipocytes were isolated from C57BL6 mice and differentiated in vitro. Once differentiated, cells were stimulated with IL-35 (100 ng/ml) for 48h in the presence or absence of SB203580 inhibitor (10 uM). The expression of principal adipogenic markers (*Pparg, Adipoq, Leptin, Perlinipin*) level was measured by qRT-PCR and relativized to *b-actin*. Data are mean ± SEM, *p < 0.05, ** p < 0.01, ***p<0.001. Analysis by *t* test (A, C, E, F), 2-way ANOVA (B) or 1-way ANOVA (G).

## DISCUSSION

Our data highlight a new role of p38 pathway in Treg cells that impairs BAT activation and browning during obesity. We have elucidated the molecular mechanism by which p38 activation regulates the production of IL35 by Treg cells through the mTOR protein pathway. Furthermore, we have uncovered a novel role of IL35 in the regulation of BAT thermogenesis. We have shown that IL-35 directly targets adipocytes and activates ATF-2, thereby stimulating the expression of UCP1 and FGF21, which in turn drive the activation of thermogenesis. In line with this, individuals with obesity exhibit lower expression of this cytokine in visceral fat. In summary, our findings reveal the novel role of IL-35 derived from Treg cells in controlling adipose tissue metabolism and function (Fig. 8).

**Fig. 8.**
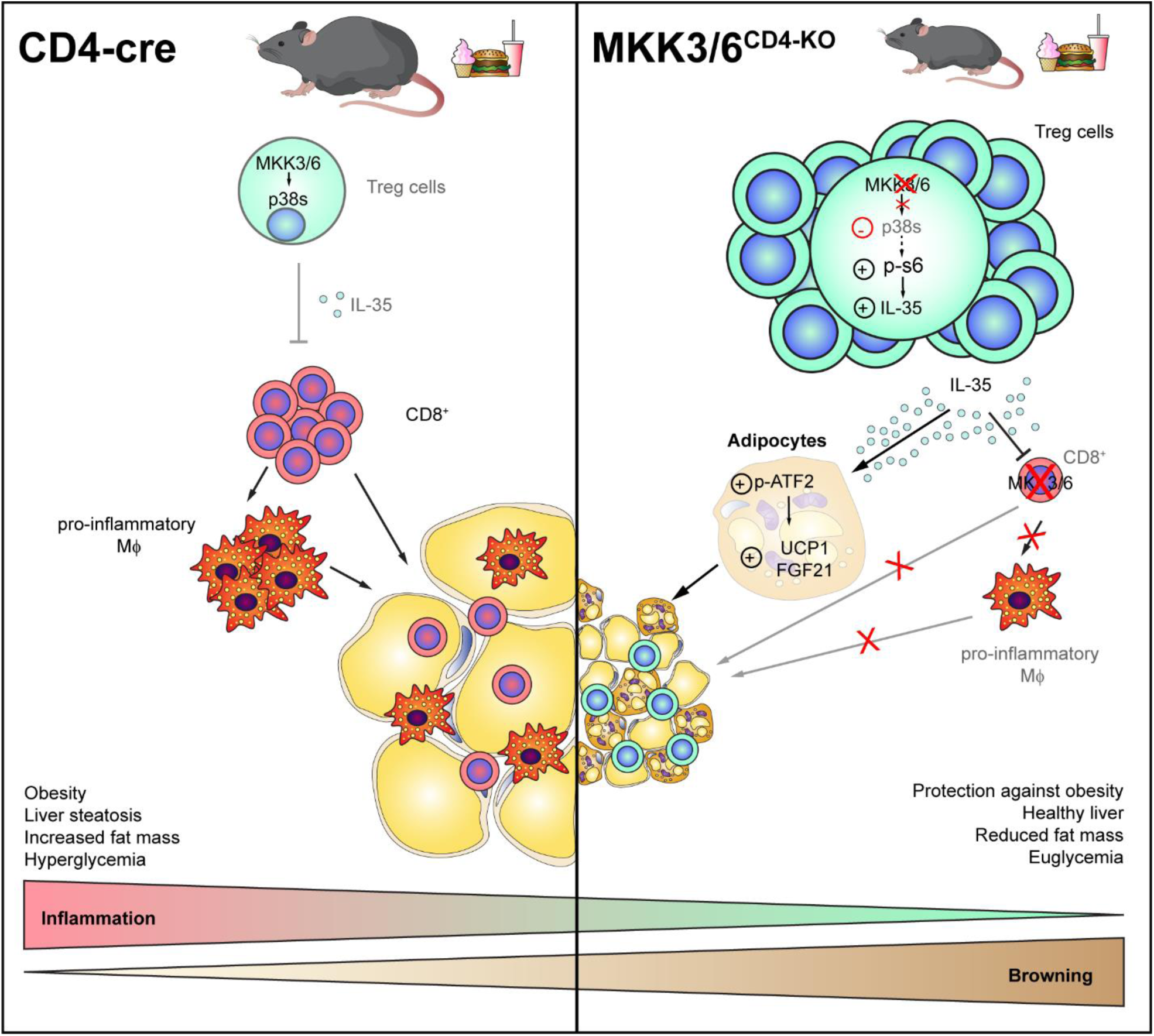
Schematic representation of how T cell p38 activation reduces Treg cells accumulation and promotes inflammation in adipose tissue via IL-35 cytokine. T cell expression of stress kinases MKK3 and MKK6 contributes to obesity and inflammation in AT through a compromised metabolic profile characterized by reduced browning and thermogenesis, as well as worsened liver steatosis. When T cells lack p38 activation, it enhances the expansion of adipose tissue Tregs and increases Treg IL35 expression via the mTOR pathway. IL35 induces phosphorylation of ATF-2, leading to the upregulation of UCP1 and FGF21 levels in adipocytes, which in turn promotes browning. Additionally, IL35 limits the infiltration and inflammation of CD8^+^ T cells in adipose tissue, thereby providing protection against obesity.

The activation of stress kinases is associated with obesity and metabolic diseases in mouse and human in a tissue specific manner (1). The importance of these kinases in myeloid cells in obesity and associated comorbidities has been widely studied. For example, p38 kinases activation polarizes macrophages towards pro-inflammatory phenotype leading to pro-inflammatory cytokine production in a model of liver steatosis (20) and LPS-induced acute hepatitis (19). In neutrophils p38 kinase family members control their migration and promotes nonalcoholic liver diseases development (3). However, our knowledge about the role of p38 kinases in T cells and their contribution in obesity development was insufficient. Analyzing the single-cell RNA sequencing data from human white adipose tissue was obtained from Emont et al. (23), we focused on the expression of p38 signalling pathway in T cells and Treg cells from adipose tissue. We observed up-regulated expression of p38α (*MAPK14)*, its activators MKK3 (*MAP2K3)* and p38 substrate *ATF-2*, while *DUSP1* phosphatase, which deactivates p38s, was down-regulated in human with obesity (BMI>30), suggesting a possible obesogenic role of p38 signalling pathway in both T cells and Treg cells. Furthermore, the results from our study confirmed p38 activation in T cells promotes obesity, as deficiency of the upstream activators of p38 kinases specifically in T cells impeded obesity development and improved their metabolic profile. Our data also indicate that p38 blockade in T cells is sufficient to blunt obesity-induced AT inflammation, thereby protecting HFD-fed MKK3/6^CD4-KO^ mice against the AT metabolic dysfunction known to be a driver of obesity comorbidities. Treg cells increase in lean individuals and are thought to prevent AT inflammation. Interestingly, we found more Treg cells in blood, lymph nodes and AT of HFD-fed MKK3/6^CD4-KO^ mice, resembling the phenotype typical for lean mice (11).

Treg cells are important immunosuppressive cells which maintenance self-tolerance, immune homeostasis and limit chronic inflammation (37–39). Observed accumulation of Treg cells in HFD-fed MKK3/6^CD4-KO^ mice restrained CD8^+^ T cells infiltration in AT and in consequence limited AT inflammation. Namely, the infiltration of CD8^+^ T cells in AT is one of the crucial steps in obesity development which precedes the activation and influx of pro-inflammatory macrophages (7). In agreement, we observed lower accumulation of CD8^+^ T cells in mice lacking p38s activation which reflected on decreased infiltration of pro-inflammatory macrophages and NK cells.

Recently, cytokine IL-35 was identified as a novel inhibitory cytokine produced by Treg cells which potentiate their suppressive activity and suppresses T cell proliferation (40). In our study, we shown that its expression was p38 dependent as chemical inhibition or genetical deficiency of p38 activation resulted in increased IL-35 expression in human and mouse Treg cells limiting AT inflammation in HFD-fed MKK3/6^CD4-KO^ mice. In addition, we found a novel mechanism by which p38 kinases regulate IL-35 expression through inhibition of the mTOR pathway.

Finally, we described a novel role of Treg derived cytokine IL-35 in adipose tissue functionality. This cytokine decreased in obese adipose tissue of human and mice. These results are in concordance with the recently described role for another member of the same IL family, IL27, which promotes adipocytes thermogenesis (41). Remarkably, lack of p38 activation in T cells improved thermogenic capacity of mice during HFD. This protection was maintained when mice were placed at isothermal conditions, confirming that observed phenotype is p38-T cell dependent. Since increased BAT thermogenesis is still present at isothermal housing conditions, suggest this effect of Treg cells/IL-35 on BAT thermogenesis is likely mediated by alternative mechanisms, which is needed to be confirmed by further studies. Our results suggest that higher expression of IL-35 in MKK3/6^CD4-KO^ mice might be responsible for their improved adaptive thermogenesis. In fact, we found that IL-35 is able to induce adipocyte thermogenic program. Mechanistically, IL-35 triggers phosphorylation of the transcription factor ATF-2 leading to increased UCP1 and FGF21 protein levels in adipocytes. While previous studies using loss of adipose Treg cells (11, 42, 43) or gain-of function Treg cells (44), did not results in profound obesity protection, our study showed that p38 signalling pathway is important in Treg cells and may modulate their function, finally leading to an upregulation of IL35 expression, which could be a contributing factor to the significant obesity protection observed in MKK3/6^CD4-KO^ mice. We believe that IL35’s effects on energy balance and thermogenesis are critical components of the observed protection against obesity in this model.

In summary, our findings unveil the importance of p38 signalling in the regulation of T cell function and identify the manipulation of this pathway is a promising therapeutic strategy for metabolic diseases. We demonstrate that p38s kinases reduces Treg cells in circulation and AT draining lymph nodes, hence leading to Treg reduction in AT and obesity development. Moreover, we show that Treg-derived IL-35 has an important role regulating AT browning via ATF-2/UCP1/FGF21 axis, hence improving metabolism and proposing this cytokine is a promising target for anti-obesity immunotherapy.

## MATERIALS AND METHODS

### Lead contact

Further information and requests for resources and reagents should be directed to and will be fulfilled by the lead contact, Guadalupe Sabio Buzo.

## EXPERIMENTAL DESIGN AND SUBJECT DETAILS

### Human visceral fat samples

For the analysis of visceral fat, the study population included 65 patients (52 adult patients with BMI >35) recruited from patients who underwent elective bariatric surgery at the University Hospital of Salamanca. Patients were excluded if they had a history of alcohol use disorders or excessive alcohol consumption (>30 g/day in men and >20 g/day in women) or had chronic hepatitis C or B. Control subjects (n = 12) were recruited among patients who underwent laparoscopic cholecystectomy for gallstone disease. The patient data is presented in table S5.

### Study approval

Human peripheral blood mononuclear cells (PBMC) were isolated from buffy coats obtained from healthy donors according to standard procedures. Buffy coats of healthy donors were received from the Blood Transfusion Center of the Comunidad de Madrid, and all donors signed their consent for the use of samples for research purposes. All procedures using primary human cells were approved by the Ethics Committee of Hospital Universitario de la Princesa. For visceral fat samples the population study was approved by the Ethics Committee of the University Hospital of Salamanca and the Carlos III (CEI PI 09_2017-v3) with the all subjects providing written informed consent to undergo visceral fat biopsy under direct vision during surgery. Data were collected on demographic information (age, sex, and ethnicity), anthropomorphic measurements (BMI), smoking and alcohol history, coexisting medical conditions, and medication use. All animal procedures conformed to EU Directive 86/609/EEC and Recommendation 2007/526/EC regarding the protection of animals used for experimental and other scientific purposes, enacted under Spanish law 1 1201/2005. The protocols are CNIC-07/18 and PROEX 215/18.

### Isolation, culture and stimulation of human peripheral blood CD4^+^ T Cells

Human peripheral blood mononuclear cells (PBMC) were collected from buffy coats from healthy donors by Ficoll density gradient separation, as described(45). CD4^+^ cells were isolated using the EasySep™ Human Naïve CD4^+^ T Cell Isolation Kit II. For *in vitro* Treg differentiation, isolated cells were activated with plate-bound anti-CD3 (3 μg/ml) and RPMI 1640 medium containing 10% FBS, 1% penicillin/streptomycin, 50 µM 2-mercaptoethanol, 2 mM L-glutamine, 1% non-essential AA, 1% anti-mycotic, 2.5 ng/ml TGF-b, and 50 U/ml IL-2 with or without 10 μM BIRB796 for 6 days in a humidified atmosphere (5% CO2, 95% air) at 37 °C.

### Mice

Floxed mutant mice for *Map2k6* (*Mkk6*) genes were as described (46). Mice with a germ-line mutation in the *Map2k3* gene and LoxP elements inserted into two introns (*Map2k3*LoxP) were generated after homologous recombination in ES cells, obtained from EUCOMM clon EPD0160_3_H09. The ES cell clones were injected into C57BL/6J blastocysts to create chimeric mice that transmitted the mutated *Map2k3* allele through the germ line. The Flp NeoR cassette was excised by crossing these mice with ACTB:FLPe B6;SJL mice, which express a FLP1 recombinase gene under the direction of the human ACTB promoter. These animals were crossed with Tg (CD4-cre)1Cwi/BfluJ mice on the C57BL/6J background (Jackson Laboratory) to generate mice lacking MKK3 and MKK6 in T cells (MKK3/6^CD4-KO^). TSC1^CD4-KO^ (Tsc1_lox (lox/lox) Tg.Cd4-Cre) were from Dr. Alejo Efeyan (CNIO). All mice were maintained on a C57BL/6J background (back-crossed for 10 generations). Genotype was confirmed by PCR analysis of genomic DNA. Mice were fed an ND or an HFD (Research Diets Inc.) for 8 weeks ad libitum. Mice were housed at temperature standard for our animal facility (23-25 °C). For thermoneutrality experiments, mice were housed at 30 °C and fed HFD for 8 weeks (until the end of the experiment).

### BAT Temperature

BAT-adjacent interscapular temperature was quantified from thermographic images captured with a FLIR T430sc Infrared Camera (FLIR Systems, Inc., Wilsonville, OR) and analysed with FLIR software.

### IL-35 administration

WT (C57BL6) mice were injected with IL-35 (300 ng/mouse, 010-001-B66, Rockland Immunochemicals, Inc.) or PBS i.v. in the retro-orbital sinus. 4 hours later BAT-adjacent interscapular temperature was measured and quantified.

### Isolation, culture and *in vitro* induction of murine Treg Cells

For *in vitro* Treg differentiation, naïve CD4^+^ T cells were isolated from the spleens of CD4-Cre, MKK3/6^CD4-KO^ or TSC1^CD4-KO^ mice using the EasySep™ Mouse Naïve CD4^+^ T Cell Isolation Kit. Cells were plated on plates previously coated with 2 μg/mL anti-CD3 antibody and incubated with 2 μg/mL anti-CD28, 2 ng/mL TGF-β1 and 20 ng/mL IL-2 in RPMI1640 supplemented with 10% FBS, 1% penicillin/streptomycin, 50 µM 2-mercaptoethanol, 2 mM L-glutamine, and 1% anti-mycotic for 96 hours in a humidified atmosphere (5% CO2, 95% air) at 37°C. At the end of the experiment, iTreg cells were collected for RNA and protein isolation or treated with rapamycin (100nm) for 4 hours and used for FACS analysis.

### Adipocytes *in vitro* differentiation and IL-35 stimulation

Immortalised brown preadipocytes from WT mice were differentiated to brown adipocytes in 10% FCS medium supplemented with 20 nM insulin, 1 nM T3, 125 μM indomethacin, 2 μg/ml dexamethasone, and 50 mM IBMX for 48 hours and maintained with 20 nM of insulin and 1 nM of T3 for 8 days. Primary white preadipocytes were isolated from C57BL/6 mice were differentiated to adipocytes for 9 days in 8% FCS medium supplemented with 5 μg/ml insulin, 25 μg/ml IBMX, 1 μg/ml dexamethasone, and 1 μM troglitazone (we changed medium every other day and supplemented with insulin and troglitazone, or the last changes was with insulin only). Differentiated adipocytes were stimulated in the presence or absence of 100 ng/ml IL-35 (010-001-B66, Rockland) for 0-2 hours or for 48 hours and then the cells were lysed for western blot. In some experiments we used 10 μM SB20253580 for 48h together with IL-35.

### Glucose measurement

Mice were starved overnight (fasted) or for 1 hour (fed) and blood glucose levels were quantified with an Ascensia Breeze 2 glucose meter.

### GTT

Overnight-starved mice were injected intraperitoneally with 1 g/kg of body weight of glucose, and blood glucose levels were quantified with an Ascensia Breeze 2 glucose meter at 0, 15, 30, 60, 90, and 120 minutes post injection.

### ITT

ITT was performed by injecting intraperitoneally 0.75 IU/kg of insulin at mice starved for 1 hour and detecting blood glucose levels with a glucometer at 0, 15, 30, 60, and 90 minutes post injection.

### Indirect calorimetry system

Energy expenditure, respiratory exchange, and food intake were quantified using the indirect calorimetry system (TSE LabMaster, TSE Systems, Germany) for 3 days.

### Nuclear magnetic resonance imaging analysis (MRI)

Body, fat, and lean mass were measured by nuclear magnetic resonance (Varian-Agilent, MA, USA) and analysed with Fiji software (Image J).

### Histology

Fresh livers, eWAT, sWAT and BAT were fixed in 10% formalin, included in paraffin, and cut into 5 μm slides followed by haematoxylin–eosin staining. Fat droplets in liver were detected by oil-red staining (0.7% in propylenglycol) in 8 μm liver slides included in OCT compound (Tissue-Tek).

### Tissue processing for flow cytometry

At the end of experiments, mouse axillar and inguinal lymph nodes, blood (100 μl), and spleens were collected, and single-cell suspensions was obtained by passing through a 70-μm cell strainer. Erythrocytes in pellets from all tissues (except lymph nodes) were lysed with a red cell lysis buffer, and the remaining cells were subsequently resuspended in flow cytometry buffer (2 PBS+1% FBS+ 2 mM EDTA). Epidydimal white adipose tissue (eWAT) was carefully excised, minced, and digested with 1 mg/mL liberase + 2 U/ml DNAse in HBSS for 25-30 min at 37 °C with shaking at 1200 rpm. Digestion was stopped by addition of PBS+10% FBS, and cells were passed through a 70-μm cell strainer and centrifuged for 5 min at 500 g to obtain the stromal-vascular fraction (SVF). SVF pellets were also used for RNA isolation and subsequent qRT-PCR analysis.

### Flow cytometry and cell sorting

Single-cell suspensions were stained with surface antibodies, and dead cells were excluded by nuclear staining with DAPI or Zombi Aqua™. Fluorochrome-conjugated antibodies to surface proteins were as follows: anti-CD45 V450 (clone 30-F11), anti-CD3 AF700 (clone 17A2), anti-CD4 APC-Cy7 (clone RM4-5), anti-CD8 BV785 (clone 53-6.7), anti-CD25 PE (clone PC61), anti-CD206 BV711 (clone C068C2), anti-F4/80 APC (clone Cl:A3-1), anti-CD11b FITC (clone M1/70), anti-CD11c PE (clone HL3), anti-IL-35 PE (clone 27537), and anti-NK1.1 PerCP-Cy5.5 (clone PK136). For intranuclear staining, cells were fixed and permeabilized with the eBioscience Foxp3 Transcription Factor Staining Buffer Kit followed by anti-Foxp3 APC (clone FJK-16s) staining. For intracellular staining of IL-35, cells were activated with PMA/ionomycin/Brefeldin A for hours before surface staining. After staining, cells were passed through 70 μm filters, and data were acquired with a BD FACSymphony flow cytometer and analysed with FlowJo software. The gating strategy is presented in figure S3A and S3B.

Single-cell suspensions of splenocytes from mice were stained with the following surface antibodies: anti-CD4 APC-Cy7 (clone RM4-5), anti-CD8 BV785 (clone 53-6.7), anti-CD3 AF700 (clone 17A2), anti-CD11b FITC (clone M1/70), and anti-CD45 V450 (clone 30-F11). Nuclei were stained with DAPI to distinguish live and dead cells. Cells were sorted with a BD FACS Aria cell sorter into the following populations: CD4^+^ T cells (CD45^+^CD3^+^CD4^+^CD8^-^), CD8^+^ T cells (CD45^+^CD3^+^CD8^+^CD4^-^) and NK cells (CD45^+^NK1.1^+^CD3^-^). Sorted cells were collected, lysed, and analysed by western blot to check MKK3 and MKK6 deletion.

### Western blot

Samples were lysed in Triton lysis buffer [20 mM Tris (pH 7.4), 1% Triton X-100, 10% glycerol, 137 mM NaCl, 2 mM EDTA, 25 mM β-glycerophosphate, 1 mM sodium orthovanadate, 1 mM phenylmethylsulphonyl fluoride, and 10 µg/mL aprotinin, and leupeptin]. Extracts (20–50 µg protein) were examined by immunoblot. Primary antibodies used in the study: anti-mouse MKK3 (Cat# 9238, Cell Signaling Technology), anti-mouse MKK6 (Cat# ADI-KAP-MA014-E, Enzo Life Sciences), anti-mouse UCP1 (Cat# AB10983, Abcam), anti-mouse FGF21 (Cat# RD281108100, BioVendor), anti-mouse p-ATF2 T69/71 (Cat# 9225S, Cell Signaling Technology), anti-mouse ATF2 (Cat# 9226S, Cell Signaling Technology), anti-mouse p-s6 S240/244 (Cat# 5364S, Cell Signaling Technology), anti-mouse p-p38 T180/Y182 (Cat# 9211S, Cell Signaling Technology), anti-mouse b-actin (Cat# sc-47778, Santa Cruz Technology), anti-mouse vinculin (Cat# V9131, Sigma) and secondary antibodies used in the study: goat anti-mouse (Cat# 31430, ThermoFisher) and goat anti-rabbit (Cat# 31460, ThermoFisher). Reactive bands were detected by chemiluminescence.

### qRT-PCR

RNA (1 μg) extracted with the RNeasy Plus Mini kit or RNeasy Plus Micro kit (Quiagen) was transcribed to cDNA, and qRT-PCR was performed using the Fast Sybr Green probe (Applied Biosystems) and the appropriate primers in a 7900 Fast Real Time thermocycler (Applied Biosystems). Data were analysed with SDS2.4 software (Applied Biosystems), and relative mRNA expression was normalized to *GAPDH* (human data) or *b-actin* (mouse data) mRNA measured in each sample. Primers used are listed in table S6.

### Single-cell RNA-Seq processing

Single-cell RNA sequencing data from human white adipose tissue was obtained from Emont et al. (23) (https://gitlab.com/rosen-lab/white-adipose-atlas). The Seurat RDS file for immune cells was downloaded and processed with Seurat 4.0 (47) (https://satijalab.org/seurat/). The clusters of T cells and Treg cells were subset and exported for further analysis. As the clusters were already identified, integration was not re-run, but we applied a linear transformation to scale the data from the T cells and Treg cells isolated cluster, followed by dimension reduction analysis (PCA). Following PCA analysis, the first 20 and 30 dimensions were used for further analysis in the T cells and Treg cells, respectively. We performed non-linear dimension reduction analysis using the UMAP algorithm. The analysis was done using the first 20 and 30 dimensions for the T cells and Treg cells clusters, respectively, and a resolution of 0.5. Differentially expressed genes between BMI ranges were identified with a non-parametric Wilcoxon rank sum test.

### Statistical analysis

Results are expressed as mean ± SEM. Statistical differences were analysed by the Student *t* test or 2-way ANOVA, with differences at *p* < 0.05 considered significant. When variances were different, ANOVA coupled with Bonferroni’s post-tests. When variances were different in *t* test, we used Welch’s test. All analyses were performed with Excel (Microsoft Corp.) and GraphPad PRISM 8 software. Statistical details for individual the experiments were indicated in the figure legends.

## Acknowledgments

We thank S Bartlett for English editing and F. Sanchez Madrid for providing human buffy coats. We thank the staff at the CNIC Cellomics and Advanced Imaging units for technical support and help with data analysis.

## Author contributions

G.S. and I.N. conceived the project, designed the study, developed the hypothesis, and generated project resources. I.N. performed the experiments, analysed the data, and prepared figures. J.A.L. and J.V. provided expertise and critical feedback. R.R.B. performed scRNA-Seq data computational analysis. A.B.P., A.E. and M.B.A. assisted in experimental analysis. M.C, A.M, L.L.V, M.L, E.R, I.R.G. and L.M. participated in the execution of experiments. L.H.C, J.L.T. and M.M. provided human visceral fat samples. P.M. and M.M. provided expertise and critical feedback. G.S and I.N. wrote the manuscript. All authors revised and approved the final manuscript.

## Data and materials availability

All data generated or analyzed during this study are included in the main text or its supplementary material. Any further information is available from the corresponding author.

## Supplementary Materials for

**Fig. S1.**
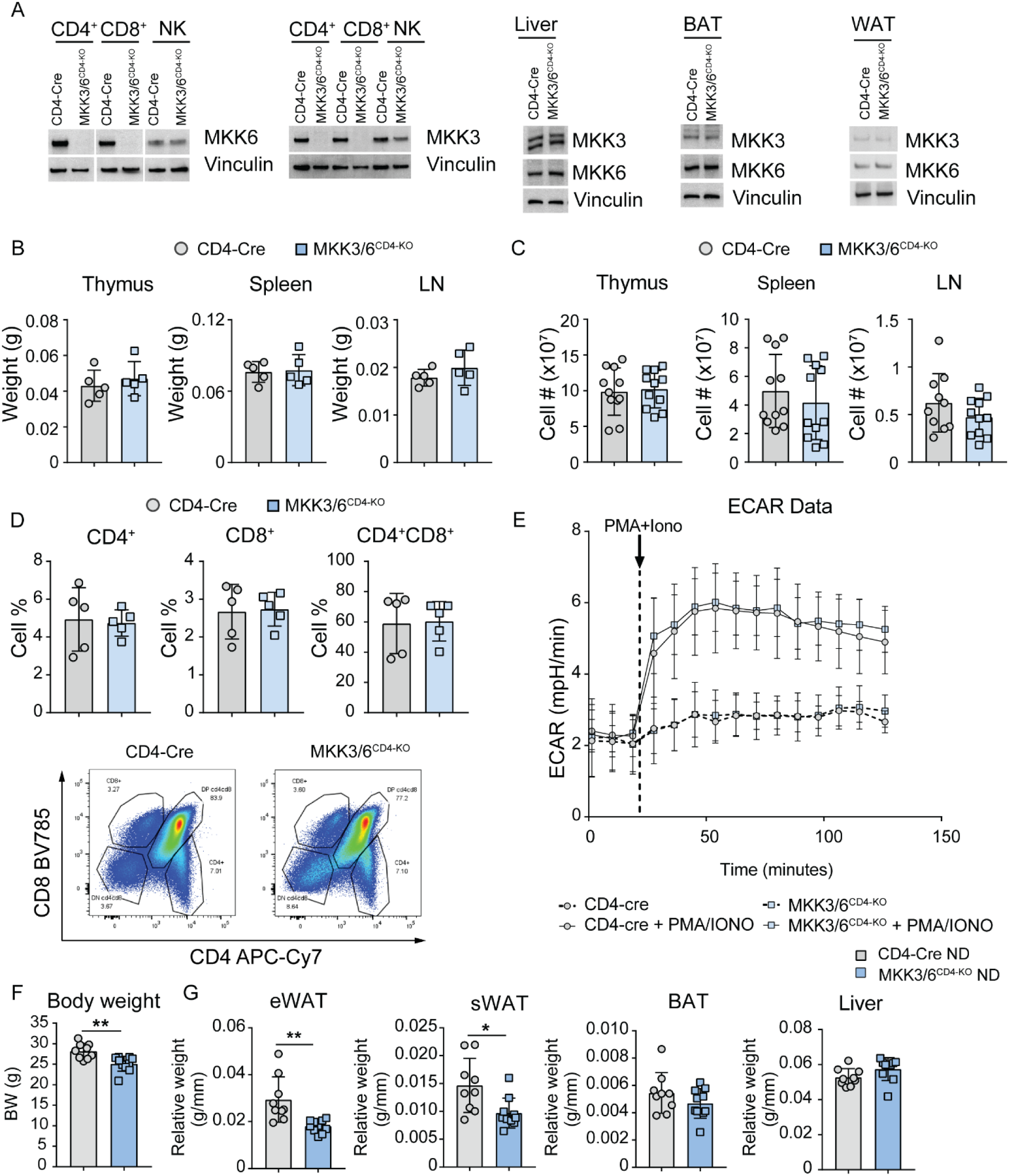
p38 MAPK activation deficiency does not induce problems in T cell development and activation. (**A**) Western blot analysis of MKK3 and MKK6 expression in CD4^+^ T cells, CD8^+^ T cells, NK cells, liver, brown adipose tissue (BAT), and white adipose tissue (WAT) isolated from MKK3/6^CD4-KO^ and control (CD4-Cre) mice. CD4^+^ T cells, CD8^+^ T cells and NK cells were sorted from spleen by FACS. (**B**) Thymus, spleen and periphery lymph nodes (LN) weight of 8-weeks old control CD4-Cre and MKK3/6^CD4-KO^ mice fed a normal diet (ND) (mean ± SEM; CD4-Cre n = 5 mice; MKK3/6^CD4-KO^ n = 5 mice). (**C**) Cell number in thymus, spleen and LN of 8-weeks old control CD4-Cre and MKK3/6^CD4-KO^ mice (mean ± SEM; CD4-Cre n = 11 mice; MKK3/6^CD4-KO^ n = 11 mice). (**D**) Frequency of CD4^+^, CD8^+^ and CD4^+^CD8^+^ T cells in thymus and representative dot plots of FACS analysis of 8-weeks old control CD4-Cre and MKK3/6^CD4-KO^ mice (mean ± SEM; CD4-Cre n = 5 mice; MKK3/6^CD4-KO^ n = 5 mice). (**E**) Sea horse analysis of real time activation of naïve CD4^+^ T cells (100.000 cells/well; n=6 wells) from CD4-Cre and MKK3/6^CD4-KO^ mice with PMA/Ionomycin (50 and 500 ng/ml respectively). (**F**) Body weight evolution in CD4-Cre and MKK3/6^CD4-KO^ male (8–10-wk-old) mice fed the ND for 10 weeks. Data are presented as the increase above initial weight (left panel), and total weight at 18 weeks of age (right panel). (**G**) eWAT, sWAT, BAT, and liver mass relative to tibia length (mean ± SEM; CD4-Cre n = 9 mice; MKK3/6^CD4-KO^ n = 10 mice). Data are mean ± SEM, exact p values are shown. 2-way ANOVA coupled with Bonferroni’s post-tests (A); or t test or Welch’s test when variances were different (A-D).

**Fig. S2.**
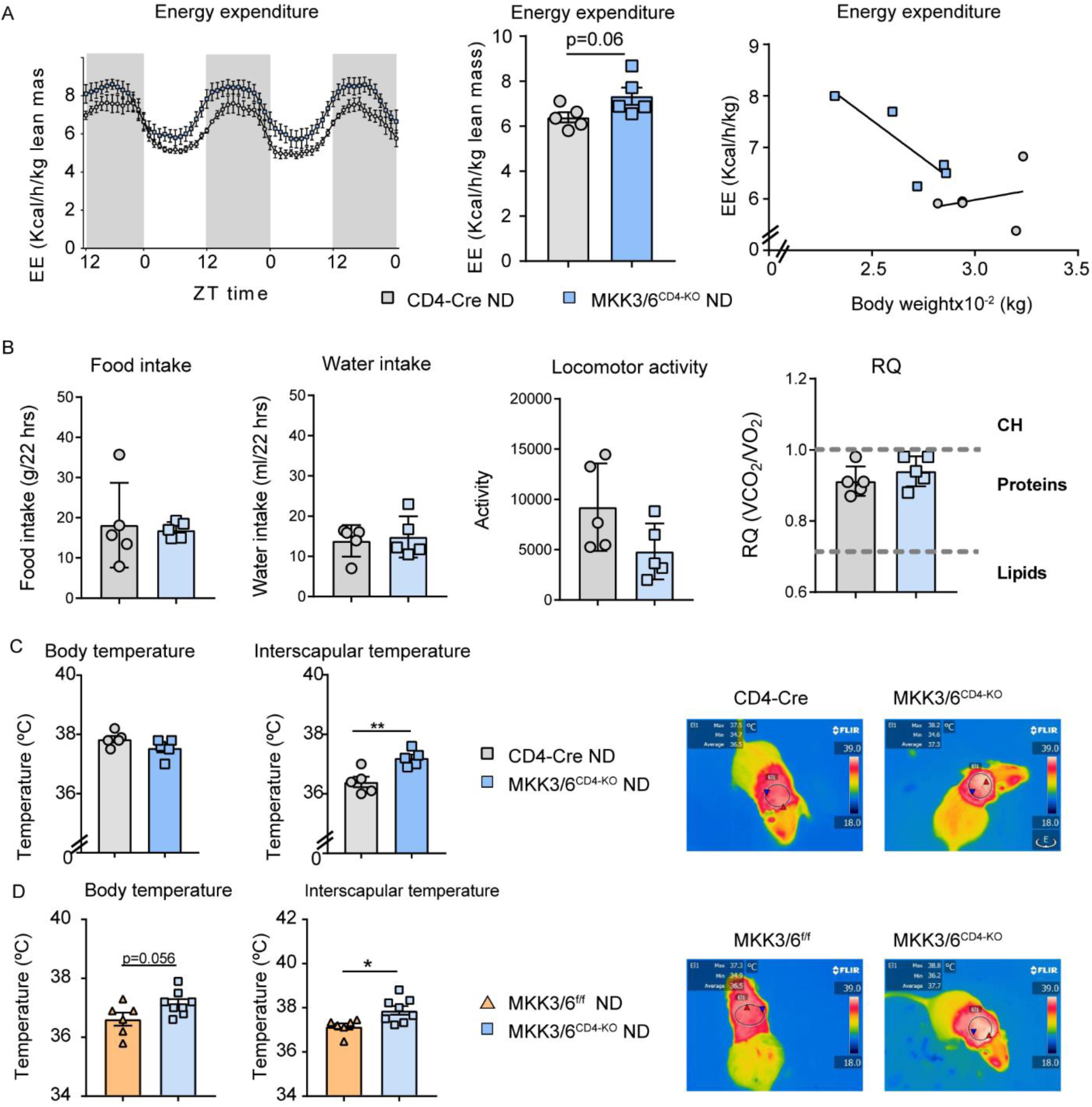
MKK3/6 deficiency in T cells increases energy expenditure and BAT temperature. (A) Comparison of energy balance between ND-fed CD4-Cre and MKK3/6CD4-KO mice examined in a metabolic cage over a 3-day period. Hour-by-hour lean–mass-corrected variation in energy expenditure (EE) (left panel; mean ± SEM; CD4-Cre n = 5 mice; MKK3/6^CD4-KO^ n = 5 mice); mean lean–mass-corrected EE (middle panel; mean ± SEM; CD4-Cre n = 5 mice; MKK3/6 ^CD4-KO^ n = 5 mice); and ANCOVA analysis of EE as a function of body weight (right panel; mean ± SEM; CD4-Cre n = 5 mice; MKK3/6 ^CD4-KO^ n = 5 mice). (B) Food and water intake, locomotor activity and respiratory quotient obtained from metabolic cages. (C) Body temperature and skin temperature surrounding interscapular BAT. Right panels show representative infrared thermal images in (C) CD4-Cre and MKK3/6 ^CD4-KO^ mice and (D) in littermates (MKK3/6^f/f^) and MKK3/6^CD4-KO^ mice fed chow diet. Data are mean ± SEM, t test *p<0.05, **p<0.01.

**Fig. S3.**
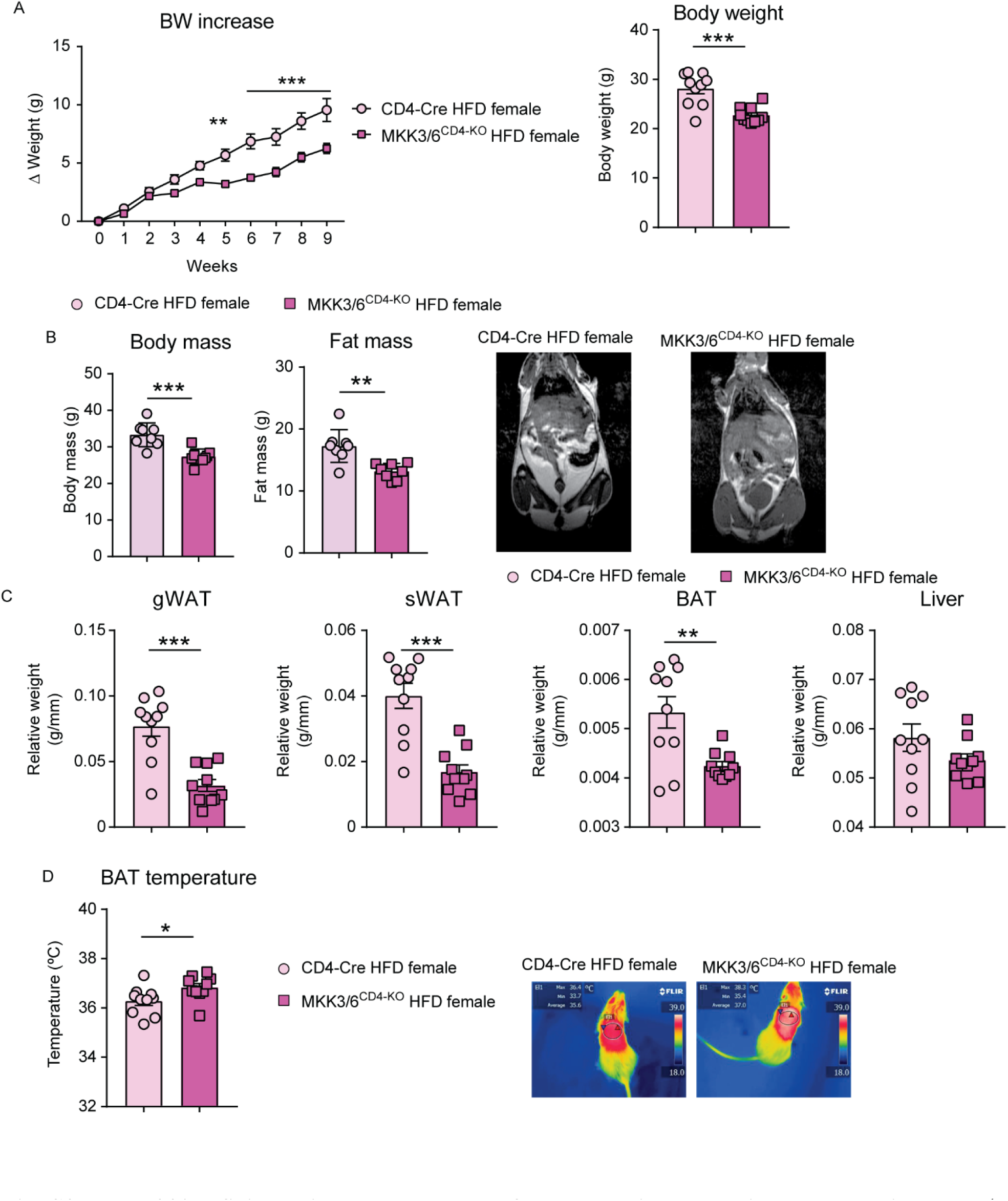
MKK3/6 deficiency in T cells protects females against HFD-induced obesity. Female MKK3/6^CD4-KO^ and CD4-Cre mice were fed a high-fat diet (HFD) for 9 weeks. (**A**) Body weight evolution in CD4-Cre and MKK3/6^CD4-KO^ female mice fed the HFD for 8 weeks (starting at 8-10 weeks old). Data are presented as the increase above initial weight (left) and absolute weight at the end of the experiment (right) (mean ± SEM; CD4-Cre n = 10 mice; MKK3/6^CD4-KO^ n = 10 mice). (**B**) NMR analysis of body and fat mass in MKK3/6^CD4-KO^ and CD4-Cre mice after 8 weeks of HFD (mean ± SEM; CD4-Cre n = 8 mice; MKK3/6^CD4-KO^ n = 9 mice). Representative images are shown on the right. (**C**) eWAT, sWAT, BAT, and liver mass relative to tibia length (mean ± SEM; CD4-Cre n = 8 mice; MKK3/6^CD4-KO^ n = 9 mice). (**D**) Skin temperature surrounding interscapular BAT. Right panels show representative infrared thermal images (mean ± SEM; CD4-Cre n = 10 mice; MKK3/6^CD4-KO^ n = 10 mice). Data are mean ± SEM, *p < 0.05, ** p < 0.01, ***p < 0.001 CD4-Cre versus MKK3/6^CD4-KO^. Analysis by 2-way ANOVA coupled to the Bonferroni post-test (A) or *t* test or by the Welch test when variances were different (A-E).

**Fig. S4.**
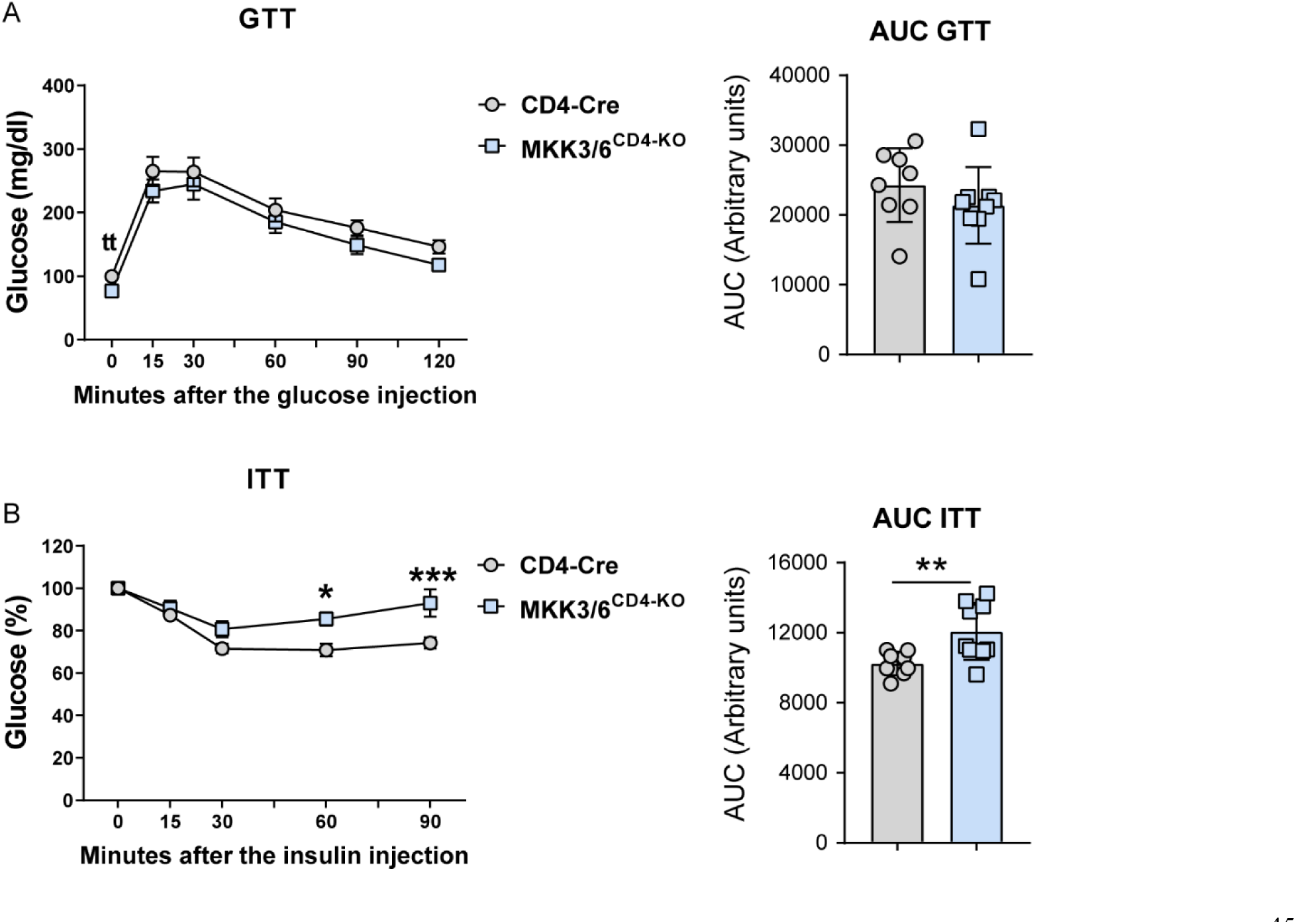
Deficiency of p38 activation in T cells slightly improves glucose tolerance and insulin sensitivity. MKK3/6^CD4-KO^ and control (CD4-Cre) mice were HFD-fed for 8 weeks. (**A**) Mice were fasted overnight (for GTT) or (**B**) 1 hour (for ITT), and blood glucose concentration was measured in mice given intraperitoneal injections of glucose (1 g/kg of total body weight) or insulin (0.75 U/kg of total body weight). Data are mean ± SEM, 2-way ANOVA *p < 0.05, ***p<0.001.

**Fig. S5.**
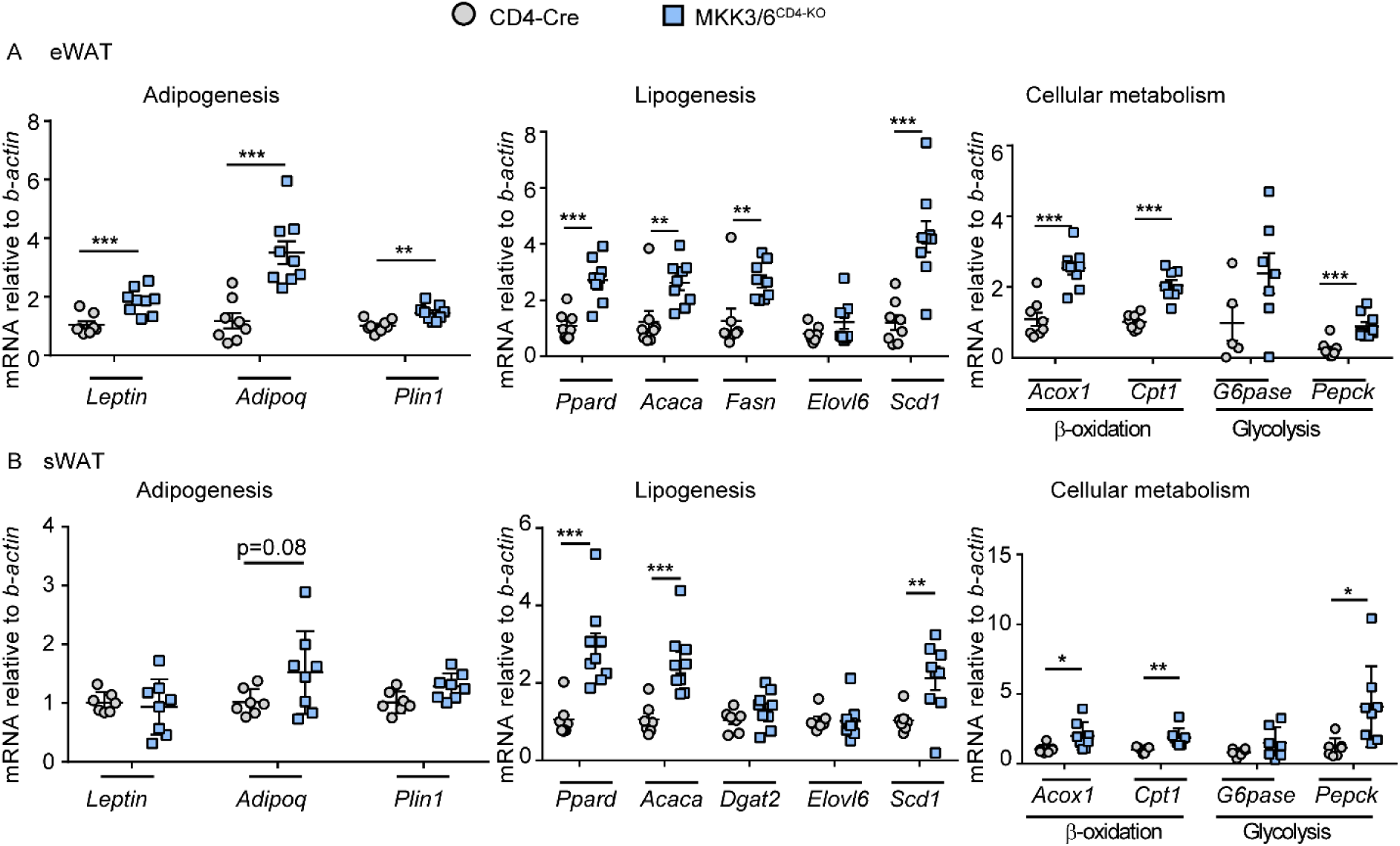
Lack of MKK3/6 in T improves AT metabolic homeostasis. MKK3/6^CD4-KO^ and control (CD4-Cre) mice were fed an HFD for 8 weeks. qRT-PCR analysis of adipogenic, lipogenic, β-oxidation, and glycolytic genes mRNA expression from (**A**) eWAT and (**B**) sWAT isolated from control or MKK3/6^CD4-KO^ mice. mRNA expression was normalized to the amount of *b-actin* mRNA. (mean ± SEM; CD4-Cre n = 8 mice; MKK3/6^CD4-KO^ n = 9 mice). Data are mean ± SEM, exact p values are shown. *t* test or Welch’s test when variances were different.

**Fig. S6.**
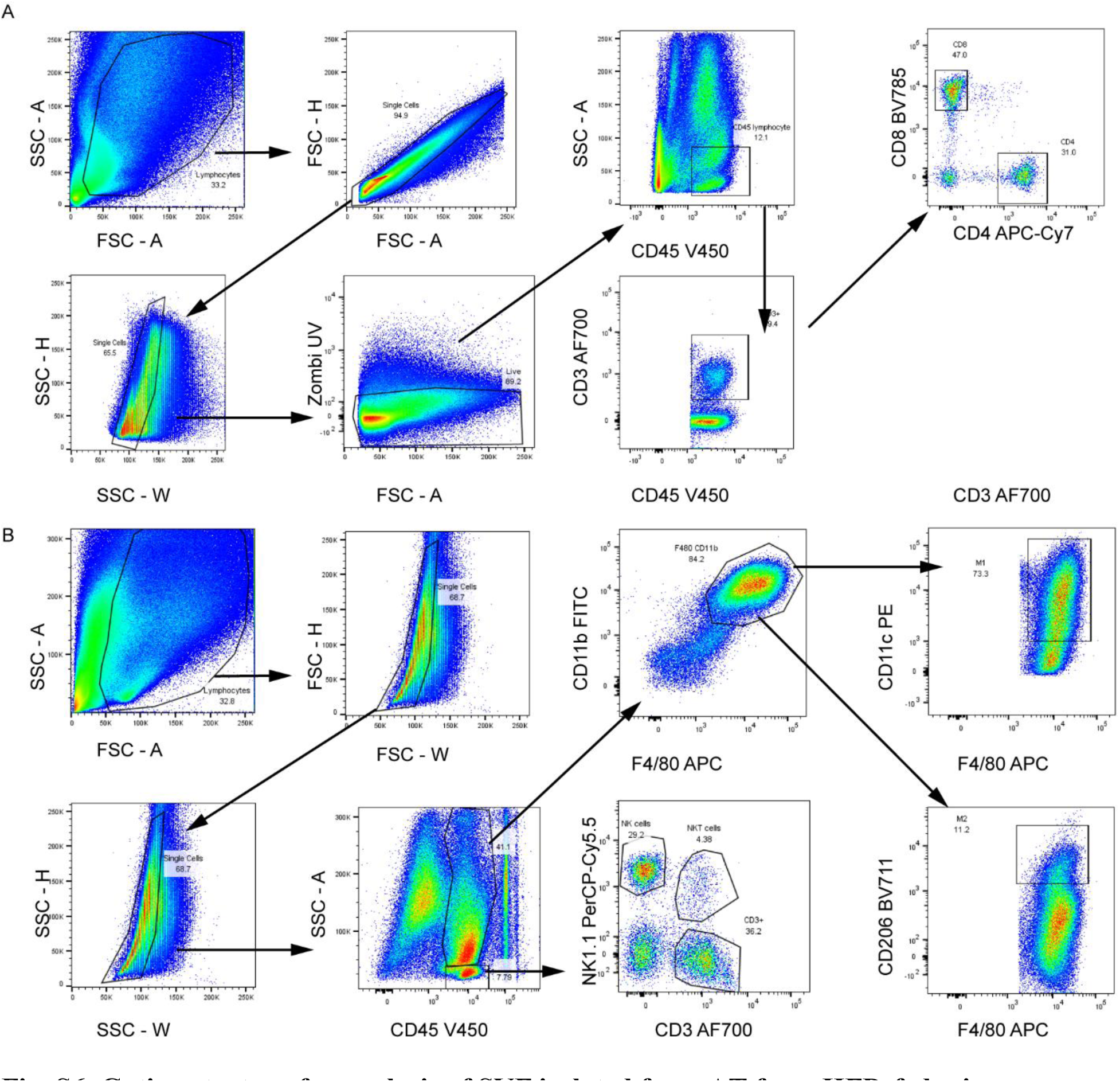
Gating strategy for analysis of SVF isolated from AT from HFD-fed mice. Representative dot plots illustrating the gating strategy for the analysis of lymphoid and myeloid populations: (**A**) CD8^+^ cells (Zombi^-^CD45^+^CD3^+^CD8^+^CD4^-^), CD4^+^ cells (Zombi^-^ CD45^+^CD3^+^CD4^+^CD8^-^); (**B**) NK cells (Dapi^-^CD45^+^NK1.1^+^CD3^-^), M1 Mϕ (Dapi^-^ CD45^+^CD11b^+^F4/80^+^CD11c^+^), and M2 Mϕ (Dapi^-^CD45^+^ CD11b^+^F4/80^+^CD206^+^). Mϕ, macrophage.

**Fig. S7.**
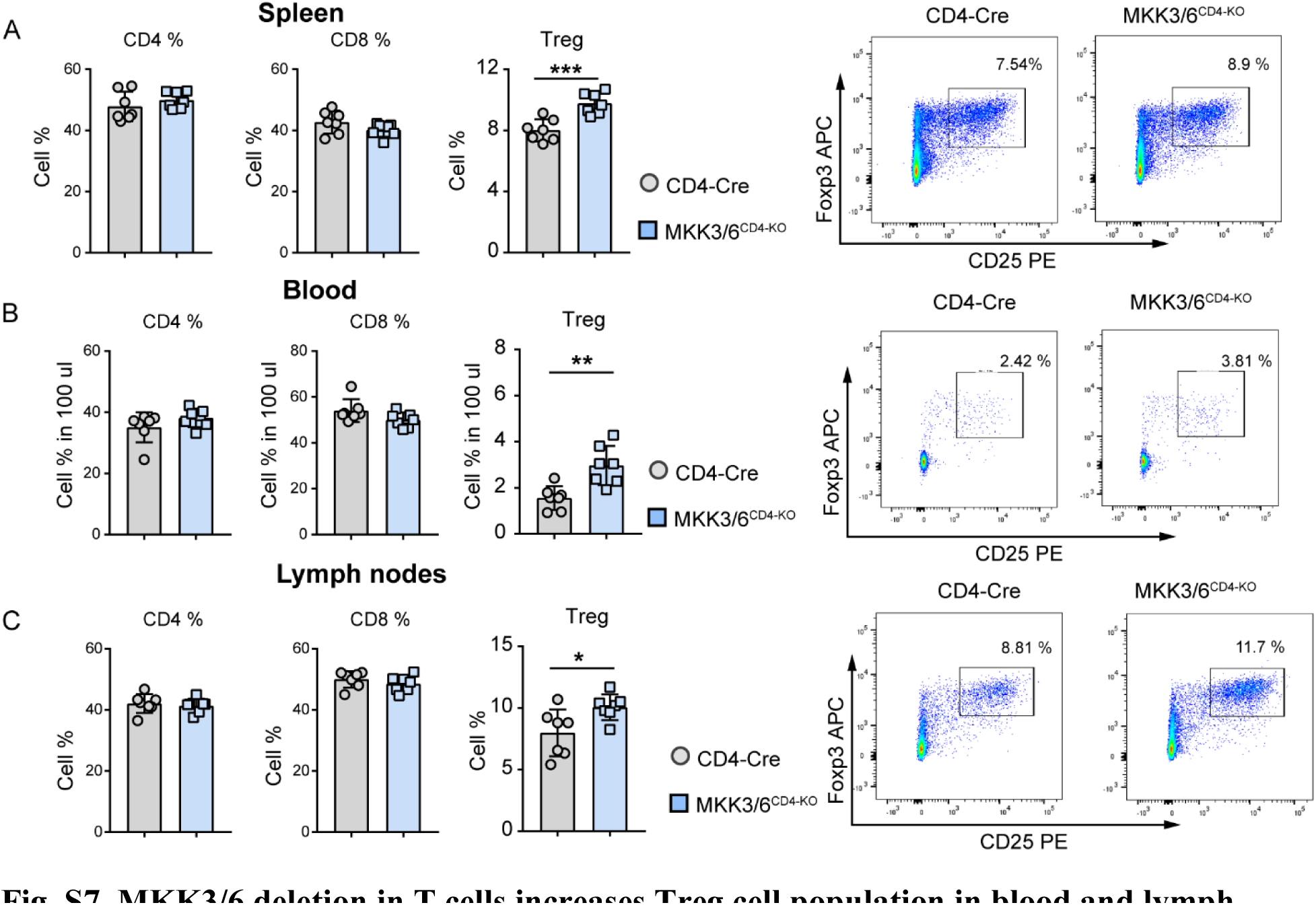
MKK3/6 deletion in T cells increases Treg cell population in blood and lymph nodes. MKK3/6^CD4-KO^ and CD4-Cre mice were fed a high-fat diet (HFD) for 8 weeks. FACS quantification and representative dot plots of CD4^+^, CD8^+^ and Treg cells (CD4^+^CD25^+^Foxp3^+^) in spleen (**A**), blood (**B**) and lymph nodes (**C**) (mean ± SEM; CD4-Cre n = 7 mice; MKK3/6^CD4-KO^ n = 7 mice).

**Fig. S8.**
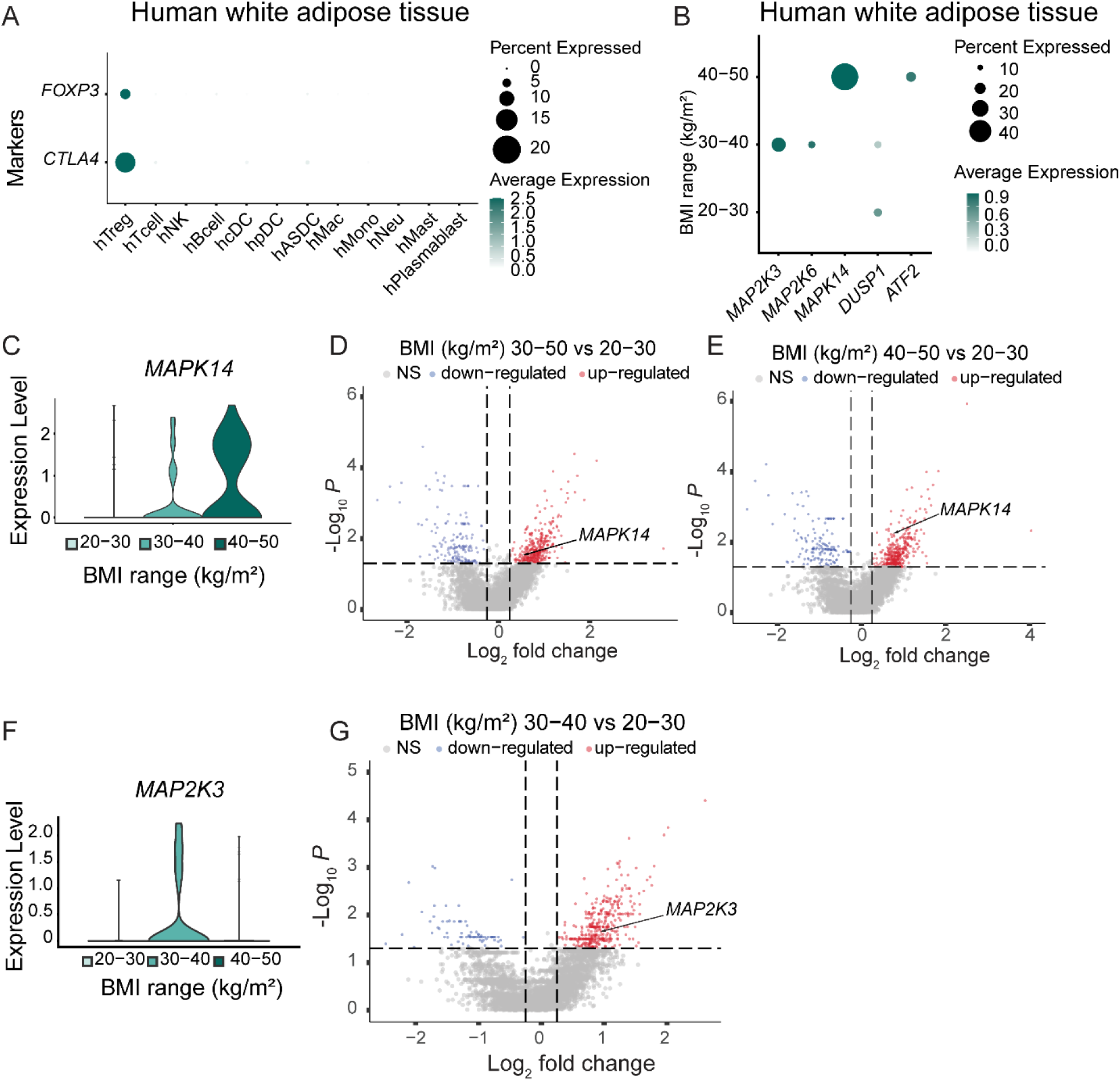
p38 MAPK pathway is upregulated in Treg cells in obese human adipose tissue. The analysis was performed using human white adipose tissue single-cell RNA-seq data from Emont et al. (23). (**A**) Dot plot of the expression of the indicated regulatory T cell (Treg) marker genes in the different cell type clusters. (**B**) Dot plot of the expression of the indicated genes by BMI range in human white adipose tissue Treg cluster shown in (**A**). (**C**) Violin plot showing the level of *MAPK14* gene expression by BMI range in the human white adipose tissue Treg dataset. (D&E) Volcano plots of differentially expressed genes in human white adipose tissue Treg subcluster in obese (BMI 30-50 kg/m^2^) versus non-obese (BMI 20-30 kg/m^2^) subjects (**D**) and in severe obese (BMI 40-50 kg/m^2^) versus non-obese (BMI 20-30 kg/m^2^) subjects (**E**). (**F**) Violin plot showing the level of *MAP2K3* gene expression by BMI range in the human white adipose tissue Treg dataset. (**G**) Volcano plots of differentially expressed genes in human white adipose tissue Treg subcluster in class 1 and 2 obesity (BMI 30-40 kg/m^2^) versus non-obese (BMI 20-30 kg/m^2^) subjects. The vertical dashed lines in D, E & G indicate a log2 fold change cut-off of 0.25. The horizontal dashed lines in D, E & G indicate a -log10 p-value cut-off of 1.3 (p-value < 0.05). DC: dendritic cells; Mac: macrophages; Mono: monocytes; Neu: neutrophiles; Mast: mastocytes.

**Table S1:** Differentially expressed genes in human white adipose tissue T cell cluster in severe obese (BMI 40-50 kg/m^2^) versus non-obese (BMI 20-30 kg/m^2^) subjects.

**Table S2:** Differentially expressed genes in human white adipose tissue Treg subcluster in obese (BMI 30-50 kg/m2) versus non-obese (BMI 20-30 kg/m2) subjects.

**Table S3:** Differentially expressed genes in human white adipose tissue Treg subcluster in severe obese (BMI 40-50 kg/m2) versus non-obese (BMI 20-30 kg/m2) subjects.

**Table S4:** Differentially expressed genes in human white adipose tissue Treg subcluster in class 1 and 2 obesity (BMI 30-40 kg/m2) versus non-obese (BMI 20-30 kg/m2) subjects.

**Table S5.** Characteristics of obese patients and controls for visceral fat analysis.

| Variable | Obese patients<br>(n = 52) | Controls<br>(n = 12) | p |
| --- | --- | --- | --- |
| BMI (kg/m<sup>2</sup>) | 49.39 (7.39) | 25.71 (3.64) | <0.0001 |
| Body fat (%) | 53.83 (4.13) | 32.26 (8.30) | <0.0001 |
| Fasting blood sugar (mg/dL) | 114.04 (47.91) | 94.73 (10.39) | 0.191 |
| Insulin (pmol/L) | 978.49 (1722.735) | 371.90 (433.92) | 0.488 |
| HOMA-IR | 8.46 (25.09) | 2.75 (3.64) | 0.478 |
| AST (IU/L) | 24.61 (12.46) | 21.45 (5.61) | 0.415 |
| ALT (IU/L) | 32.60 (18.29) | 24.73 (11.82) | 0.178 |
| Total cholesterol (mg/dL) | 193.80 (39.05) | 199.5 (43.82) | 0.683 |
| Triglycerides (mg/dL) | 156.24 (88.95) | 147.39 (56.12) | 0.765 |
| LDL-cholesterol (mg/dL) | 113.81 (38.126) | 121 (35.366) | 0.604 |
| HDL-cholesterol (mg/dL) | 47.12 (13.11) | 48.61 (18.84) | 0.773 |

Variables are presented as mean (standard deviation) and analysed by T-test. BMI: body mass index. AST: aspartate aminotransferase. ALT: alanine aminotransferase.

**Table S6.**
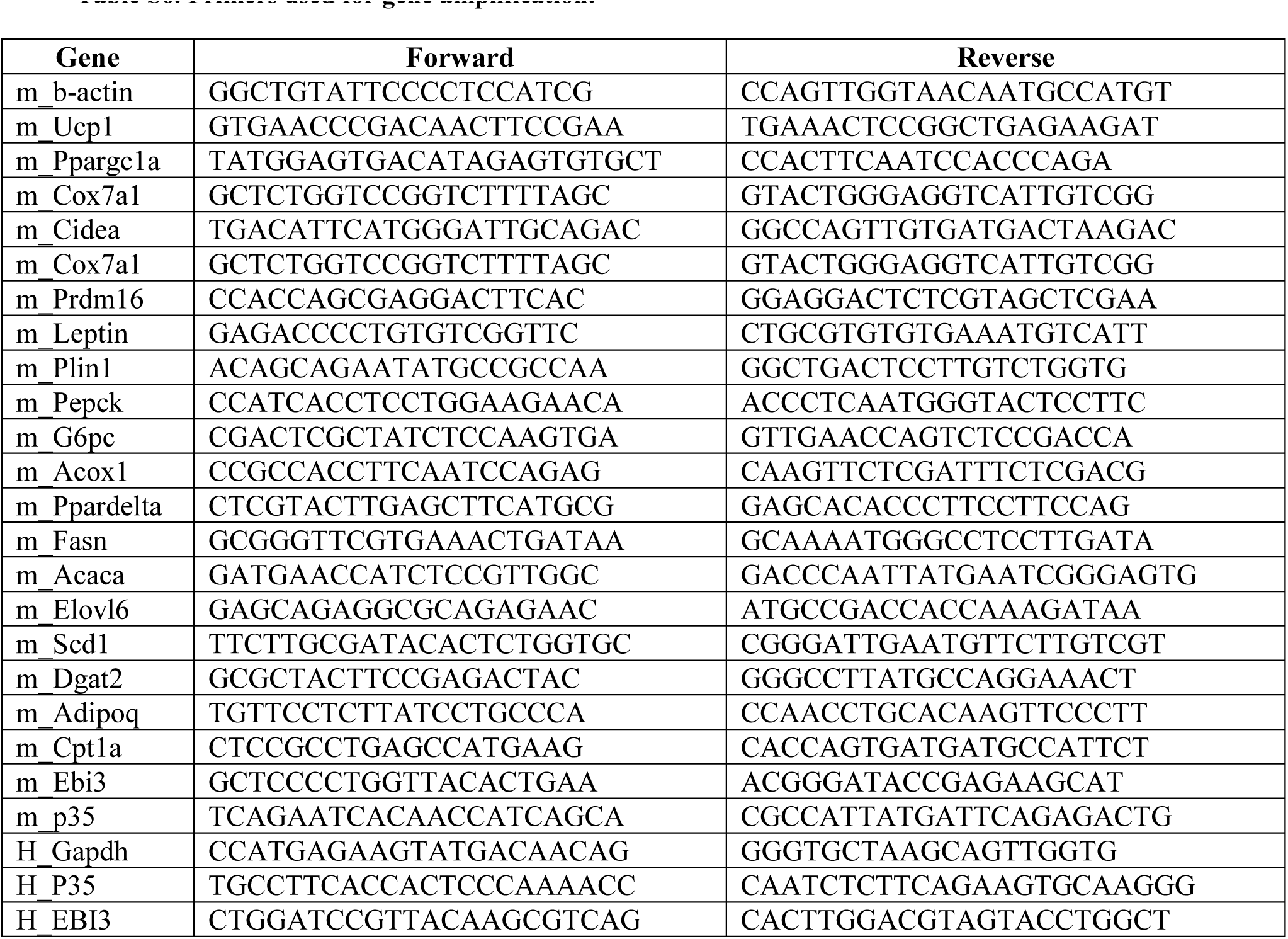
Primers used for gene amplification.

